# The establishment of *Populus* x *Laccaria bicolor* ectomycorrhiza requires the inactivation of MYC2 coordinated defense response with a key role for root terpene synthases

**DOI:** 10.1101/2022.09.06.505662

**Authors:** José Eduardo Marqués-Gálvez, Veronica Basso, Annegret Kohler, Kerrie Barry, Keykhosrow Keymanesh, Jenifer Johnson, Vasanth Singan, Igor V. Grigoriev, Rytas Vilgalys, Francis Martin, Claire Veneault-Fourrey

## Abstract

The jasmonic acid (JA) signaling pathway plays an important role in the establishment of the ectomycorrhizal symbiosis (ECM) between *Laccaria bicolor* and poplar. We previously showed that the *L. bicolor* effector MiSSP7 induces the stabilization of the poplar JAZ6, a JA co-repressor protein that binds to *Populus* MYC2.1 and MYC2.2, orthologs of the *Arabidopsis* MYC2 transcription factor (TF), blocking their activity. Here we showed that both TFs play a central role in root colonization by *L. bicolor* mycelium, since their overexpression decreased the formation of the Hartig net, the hyphal network involved in symbiotic nutrient exchanges. By combining RNA sequencing and DNA Affinity Purification sequencing (DAP-seq) analysis, we identified a core set of JA-responsive genes directly activated by poplar MYC2.1 and MYC2.2, that need to be bypassed by the fungi to colonize root apoplastic spaces. These genes encode for other TFs, receptor-like kinases and many defense-related proteins, including terpene synthases (TPS). Monoterpenes produced by some of these TPS impact *L. bicolor* growth and ECM formation, suggesting a role for poplar root monoterpenes as negative regulators of *in planta* fungal growth and ECM symbiosis.

**Significance statement:** The ectomycorrhizal symbiosis is a predominant mutualistic plant-fungus interaction occurring in forests, sustaining tree health. Ectomycorrhizal fungi colonize the root intercellularly establishing the symbiotic interface required for bidirectional nutrients exchanges, the Hartig net. During root colonization, the fungus *L. bicolor* produces the effector protein MiSSP7 that binds to the jasmonate co-receptor PtJAZ6, maintaining the repression of MYC2-targeted genes. Here we showed that defensive genes are major targets of MYC2, suggesting that their strict control is required to allow fungal colonization, with special emphasis on the host root monoterpene synthesis. Future research will focus on how root terpene defenses mediate belowground mutualistic interactions and how they can be manipulated to engineer plants with enhanced disease resistance but stable mutualistic interactions.

## Introduction

Plant roots host a wide range of microbes, including detrimental, commensal or beneficial bacteria and fungi (1). In order to interact with these micro-organisms, plants have developed recognition and defense systems that are coordinated by plant hormones, such as jasmonic acid (JA) or ethylene (ET). These hormonal pathways, and their crosstalk, are keys for the output of plant-microbe interactions, both pathogenic and beneficial (2). In particular, multiple roles have been attributed to JA, including defense against necrotrophic microorganisms and herbivores, regulation of mutualistic interactions, response to abiotic stresses, reproduction and development (3-7).

The molecular events following JA perception in plants were identified more than a decade ago (8-15). Briefly, the bioactive form JA-isoleucine (JA-Ile) is synthesized from galactolipids both in leaves and roots after wounding and perceived by the F-box protein CORONATINE-INSENSITIVE1 (COI1), triggering the ubiquitination and degradation of JASMONATE ZIM DOMAIN (JAZ) proteins. This releases the repression of different transcription factors (TFs) interacting with JAZ protein, such as the basic Helix-Loop-Helix (bHLH) TFs MYC2, MYC3, MYC4 or MYC5, which crucially regulate JA-signaling pathways by activating different sets of JA-responsive genes (16-19). MYC2 was the first of those master regulators to be characterized (20) and was shown to perform key functions related to defense reactions, including the cross-talk with other hormonal pathways (21-23), modulation of growth-defense trade-off (24) and coordinating induced systemic resistance caused by beneficial microorganisms (25). Additionally, MYC2 regulates the biosynthesis of several secondary metabolites with antiherbivore and antimicrobial activity, such as glucosinolates, threonine deaminase and proteinase inhibitor 1, steroidal glycoalkaloids and terpenes (26-31). On the other hand, JASMONATE ASSOCIATED MYC2-LIKE (JAM) proteins are also bHLH TFs, but they are positively regulated in a COI1- and MYC2-dependent manner in *Arabidopsis* and negatively regulate JA responses, mainly acting as antagonists to MYC2 (32).

Ectomycorrhizal (ECM) fungi are the most predominant belowground fungi interacting with forest trees. They form mutualistic symbiosis with roots, improving nutrient acquisition, biotic and abiotic resistance and tree intercommunication (33). During this symbiotic interaction, ECM hyphae aggregate on the surface of roots to form a mantle and then massively colonize the root apoplastic space forming a symbiotic interface, known as the Hartig net, where the exchange of nutrients takes place. ECM development entails the secretion of both fungal and plant effector proteins that play a key role in the molecular dialogue taking place between ECM symbionts (34). The first fungal effector that was found to play a central role in ECM establishment was *Laccaria bicolor* Mycorrhiza-induced Small Secreted Peptide 7 (LbMiSSP7). This effector facilitates the fungal colonization of the apoplastic space between poplar root cells that is needed to form the Hartig net (HN) (35-36). LbMiSSP7 interacts with PtJAZ6 in the nuclei of poplar root cells, preventing PtJAZ6 degradation. This enhances the interaction between PtJAZ6 and both poplar transcriptional homologs of *Arabidospsis* MYC2: PtMYC2.1 and PtMYC2.2, in particular with PtMYC2.1 but does not impact PtJAZ6-PtJAM1.1 interaction (37). MYC2 controlled-genes that need to be repressed in poplar roots for ECM establishment remain unknown so far. In this work, we showed that the overexpression of poplar MYC2 TFs results in the impairment of symbiosis establishment with *L. bicolor* coupled with a coordinated upregulation of defensive genes. Moreover, we found that both *Populus tremula x alba* PtaMYC2.1 and PtaMYC2.2 directly control the expression of a subset of JA-responsive gene candidates playing a role in the inhibition of fungal colonization. Among those, terpene synthases (TPSs) are tightly regulated by MYC2 and some of TPS volatile products impair *in vitro* fungal growth and ECM formation.

## Results

### Overexpression of *PtaMYC2*.*1* and *PtaMYC2*.*2* impairs *in planta* fungal colonization

To investigate the roles of the transcriptional activator PtaMYC2 and repressor PtaJAM1 in symbiosis development, we generated transgenic poplars overexpressing the corresponding genes. According to gene expression levels, we selected two GUS lines as controls, two MYC2.1 overexpressing (OE) lines (MYC2.1 OE2 and MYC2.1 OE3), one MYC2.2 OE line (MYC2.2 OE2) and one JAM1.1 OE line (JAM1.1 OE3) (Fig. S1A). OE lines were similar to control lines in terms of shoot weight but some differences were found in anthocyanin and chlorophyll contents in MYC2 OE lines with respect to the control (Fig. S2). Overall, measurements of root architecture features suggested that the overexpression of *PtaMYC2*.*1, PtaMYC2*.*2* and *PtaJAM1*.*1* differentially affected the root architecture, especially the number of adventitious roots (Fig. S3). Neither the number of ECM root tips (Fig. 1A) nor their diameter (Fig. S4) were different in OE lines compared to the control GUS lines. However, we found an altered HN phenotype for JAM1.1 OE3, with an increased frequency of this structure and a trend towards deeper *in planta* colonization (Fig. 1C, D). A fully defective HN phenotype was found for MYC2.1 OE3 and MYC2.2 OE2, with an almost total absence of HN accompanied by the lack of the fungal mantle only for the MYC2.1 OE3 line (Fig. 1B, C, D). Finally, a partial HN-defective phenotype was observed for MYC2.1 OE2, with a significant decrease in HN frequency but not in HN depth or mantle thickness (Fig. 1B, C, D). It is worth noting the significant lower HN frequency in MYC2.1 OE3 compared to MYC2.1 OE2, that could be related to a > 2-fold higher overexpression of *PtaMYC2*.*1* in MYC2.1 OE3 than in MYC2.1 OE2, as observed both in RT-qPCR (Fig. S1A) and in RNA-seq data (Fig. S1B). To assess whether the localization pattern of *PtaMYC2*.*1* and *PtaMYC2*.*2* expression was similar, we generated composite poplars expressing in their roots the promoter region of either *PtaMYC2*.*1* and *PtaMYC2*.*2* genes fused to GFP protein (Fig. S5). Both genes were expressed in poplar lateral root tips (Fig. S6).

**Figure 1.**
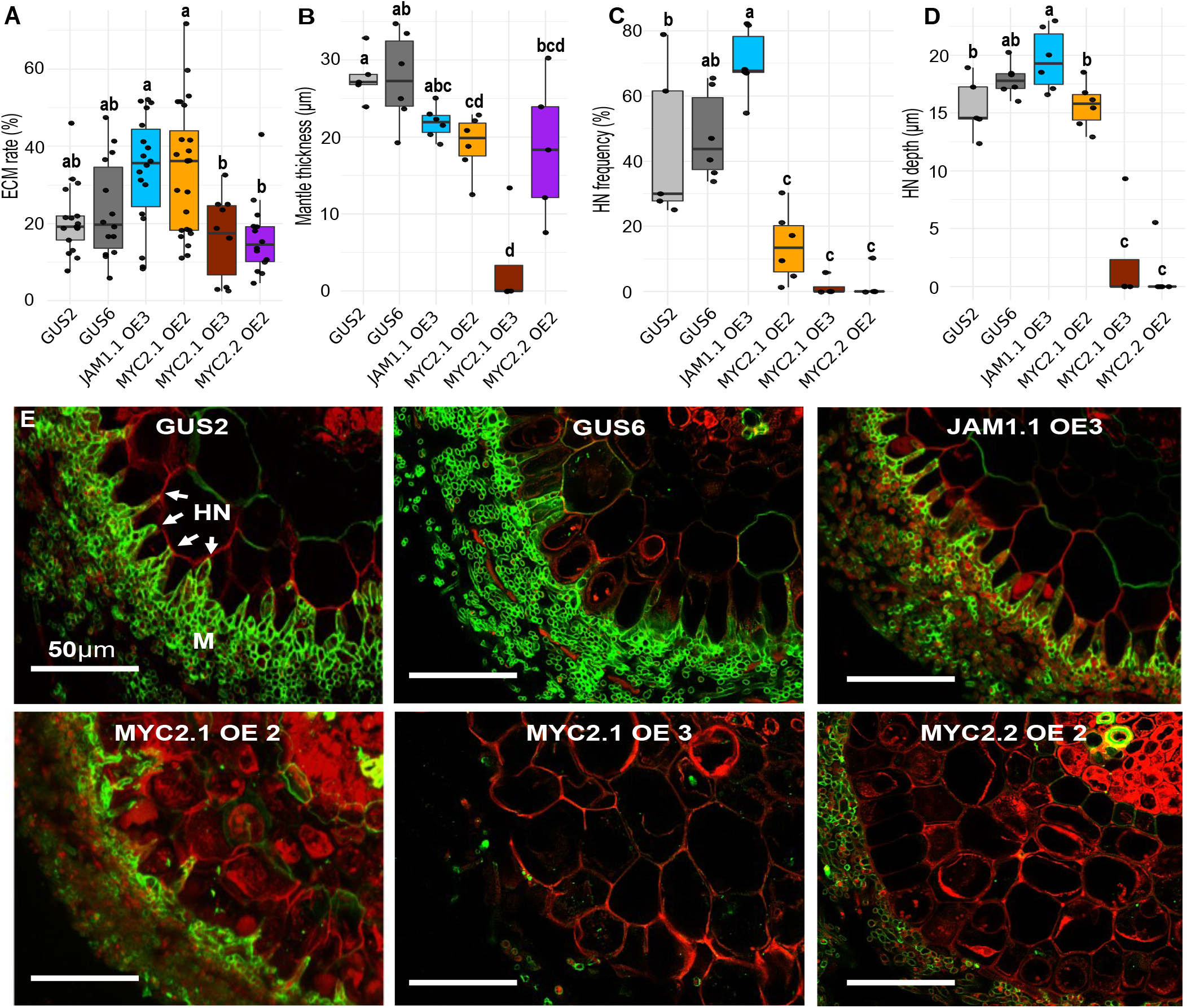
Overexpression of *PtaJAM1*.*1, PtaMYC2*.*1* and *PtaMYC2*.*2* alters ectomycorrhiza (ECM) phenotype. Boxplots represent **(A)** the ratio between ECM lateral root tips and total lateral root tips (ECM rate in percentage), **(B)** mantle thickness, **(C)** the ratio between the surface of the apoplastic space colonized by *L. bicolor* and the total apoplastic space surface (HN frequency) and **(D)** the depth of HN ingress. Whiskers represent the limits of the 1.5 interquartile range. Different letters represent significant differences based on Kruskal-Wallis one-way analysis of variance and Least Significant Difference (LSD) post-hoc test (p < 0.01), n=18 (A), n=6 (B-D). **(E)** Representative confocal microscopy images of transversal sections of each transgenic line in contact with *L. bicolor* at 21 dpi. Propidium Iodide (red) stains the root cell wall, while AlexaFluor® WGA-288 (green) stains the fungal cell-wall. Scale bar: 50 μm.

### Impairment of Hartig net formation is linked to MYC2-transcriptional core set of genes mainly consisting of defense genes

To point out differentially expressed genes (DEGs) related to altered ECM phenotypes, we compared transcripts from each transgenic line with those of GUS control lines (according to our experimental set up, see Materials and methods) (Fig. 2A, Dataset S1-S8). Several GO terms related to plant defense were significantly enriched in JAM1.1 OE3-downregulated genes both in uncolonized and colonized conditions, including those coding for chitinases, endopeptidase inhibitors, peroxidases, and terpene synthases (TPS) (Fig. S7).

**Figure 2.**
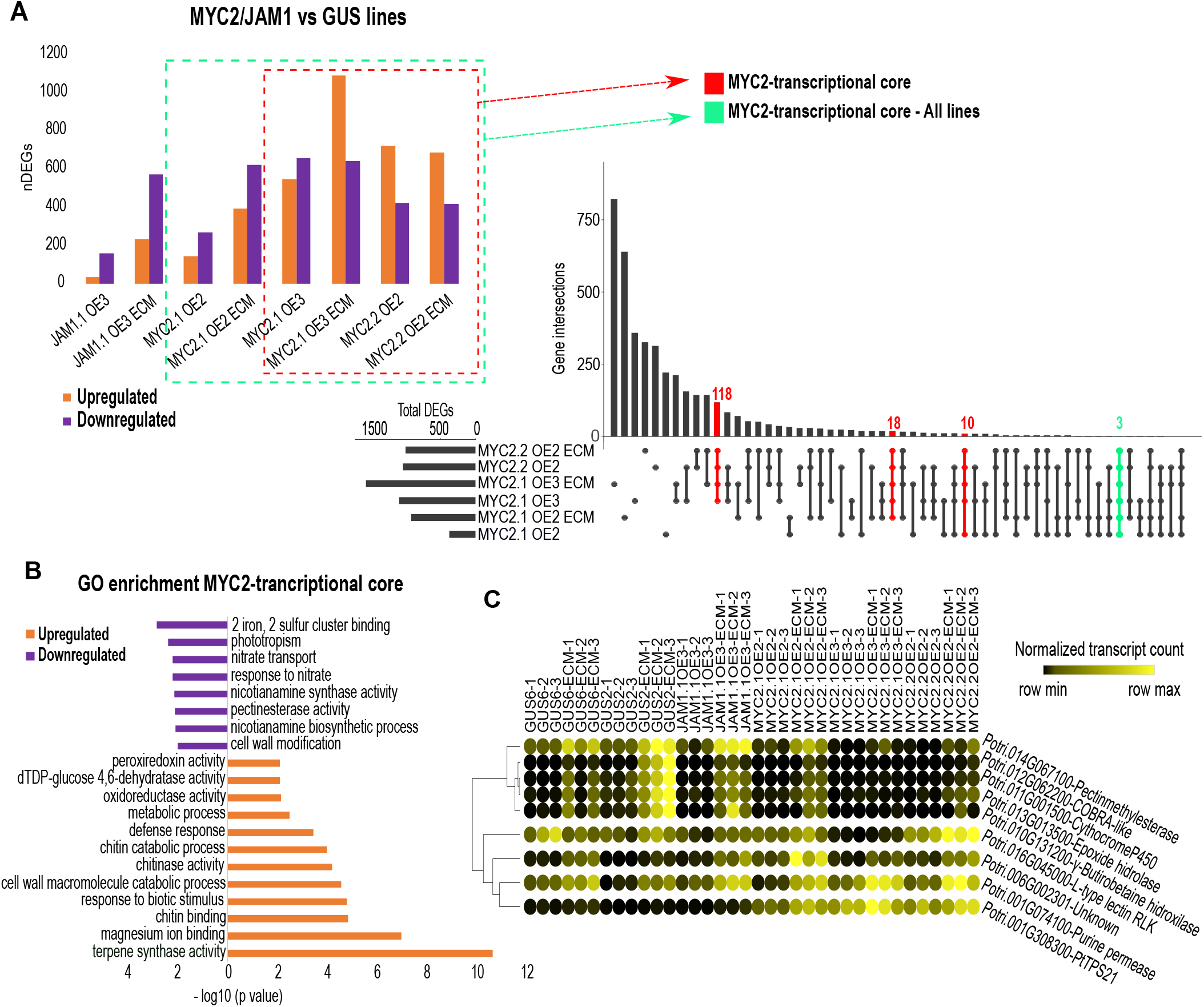
RNA-seq analysis from each *Populus tremula x alba* transgenic line reveals a core set of genes regulated by MYC2, mostly comprising upregulated defense-related genes. **(A)** The total number of down- and upregulated genes in different pairwise comparisons is shown. JAM1.1 OE3, MYC2.1 OE2, MYC2.1 OE3 and MYC2.2 OE2 lines were compared against GUS2 and GUS6 control lines, in both non-inoculated or 21 dpi (ECM) conditions (MYC2/JAM1 vs GUS lines). The MYC2-transcriptional core gene set is composed of DEGs common to MYC2 OE lines when each transgenic line was compared to GUS control lines. **(B)** GO enrichment for up- and downregulated genes (analyzed separately) belonging to the “MYC2-transcriptional core gene set”. Only significantly enriched (p < 0.01) terms are shown. **(C)** DESeq2 normalized transcript counts across all samples of those genes overlapping between the ECM-responsive gene set and the MYC2-transcriptional core gene set.

To highlight poplar genes putatively responsible of the HN-defective phenotype, we first focused on DEGs common to all transgenic lines that presented decreased HN frequency, namely MYC2.1 OE2, MYC2.1 OE3 and MYC2.2 OE2 (Fig. 2A). Only three upregulated genes were differentially expressed in all MYC2 OE lines, both in non-inoculated and ECM conditions, including two TPS-encoding genes (Potri.001G308200 and Potri.001G308300). These TPS genes have previously been characterized in *P. trichocarpa* as *PtTPS16* and *PtTPS21* (38). Another set of 146 genes were commonly regulated only in the lines presenting a complete HN-defective phenotype (MYC2.1 OE3 and MYC2.2 OE2), although 28 of them were also found to be regulated in the partial HN-defective MYC2.1 OE2 line in either ECM or uncolonized conditions. We selected these 146 plus the three common genes to all MYC2 OE lines for further analysis and named them “MYC2-transcriptional core” (Dataset S9). This core set of genes contained 59 significantly downregulated genes, with enriched functions related to nitrate transport and metabolism or plant cell wall remodeling (Fig. 2B). The other 90 DEGs were upregulated and showed significant enrichment in functions related to biotic perception and defense, similarly to those found to be downregulated in the JAM1.1 OE3 line (Fig. S8), such as peroxiredoxin, chitinase and TPS (Fig. 2B). Only nine MYC2-transcriptional core genes were also found to be differentially expressed in response to fungal colonization (ECM responsive genes, Fig. S8, Dataset S10-S16). *PtTPS21* was slightly induced upon fungal inoculation (log2 FC 1.62 ± 0.11), but its expression was > 10-fold higher in MYC2 OE than in JAM1.1 OE and GUS lines (log2 FC 4.99 ± 0.79) (Fig. 2E). The pectinmethylesterase, COBRA-like, Epoxide hydrolase, CytP450 and γ-butirobetaine hidroxilase genes clustered together according to their normalized transcript level, presenting a similar expression pattern, i.e., they were highly induced upon fungal inoculation in the HN-forming lines GUS2, GUS6 and JAM1.1 OE3, but repressed in HN-defective MYC2 OE lines, both in non-inoculated and inoculated conditions, suggesting their role as positive regulators of ECM symbiosis (Fig. 2E).

### PtaMYC2.1 and PtaMYC2.2 directly target other transcription factors and defense genes related to Hartig net inhibition

To find PtaMYC2.1 and PtaMYC2.2 direct targets, we performed DNA affinity purification sequencing (DAP-seq) experiments with these two TFs. We did not find the DNA Binding Sites (DBSs) of PtaJAM1.1 because this protein could not be correctly expressed coupled to the HALO tag *in vitro* (Fig. S9). We identified a total of 6,058 significant peaks for PtaMYC2.1 and 7,863 for PtaMYC2.2 (q < 0.1, p < 0.01) (Dataset S17-S18). In both cases, more than 50% of the peaks matched with one or several G-box-like motif (5’ CAC[C/T][A/G]TG 3’). The genes associated to these motifs are related to JA and wounding responses together with responses to environment and other stimuli, such as water deprivation, ABA or auxin (Fig. 3A). Out of the total PtaMYC2.1 and PtaMYC2.2 peaks, 1,548 (25.55%) and 1,966 (25.01%) were located in promoter regions (defined as ≤ 3 kb from the Transcription Starting Site, TSS), respectively (Fig. S10). The two TFs shared 1,104 DBSs, representing 68.70% - for PtaMYC2.1 – and 53.62% - for PtaMYC2.2 -, of their promoter DBSs (Fig. 3B). The GO enrichment test performed on promoter DBSs showed enriched functions related mainly to DNA binding and transcriptional regulation for both common and unique PtaMYC2.1 and PtaMYC2.2 sites (Dataset S19).

**Figure 3.**
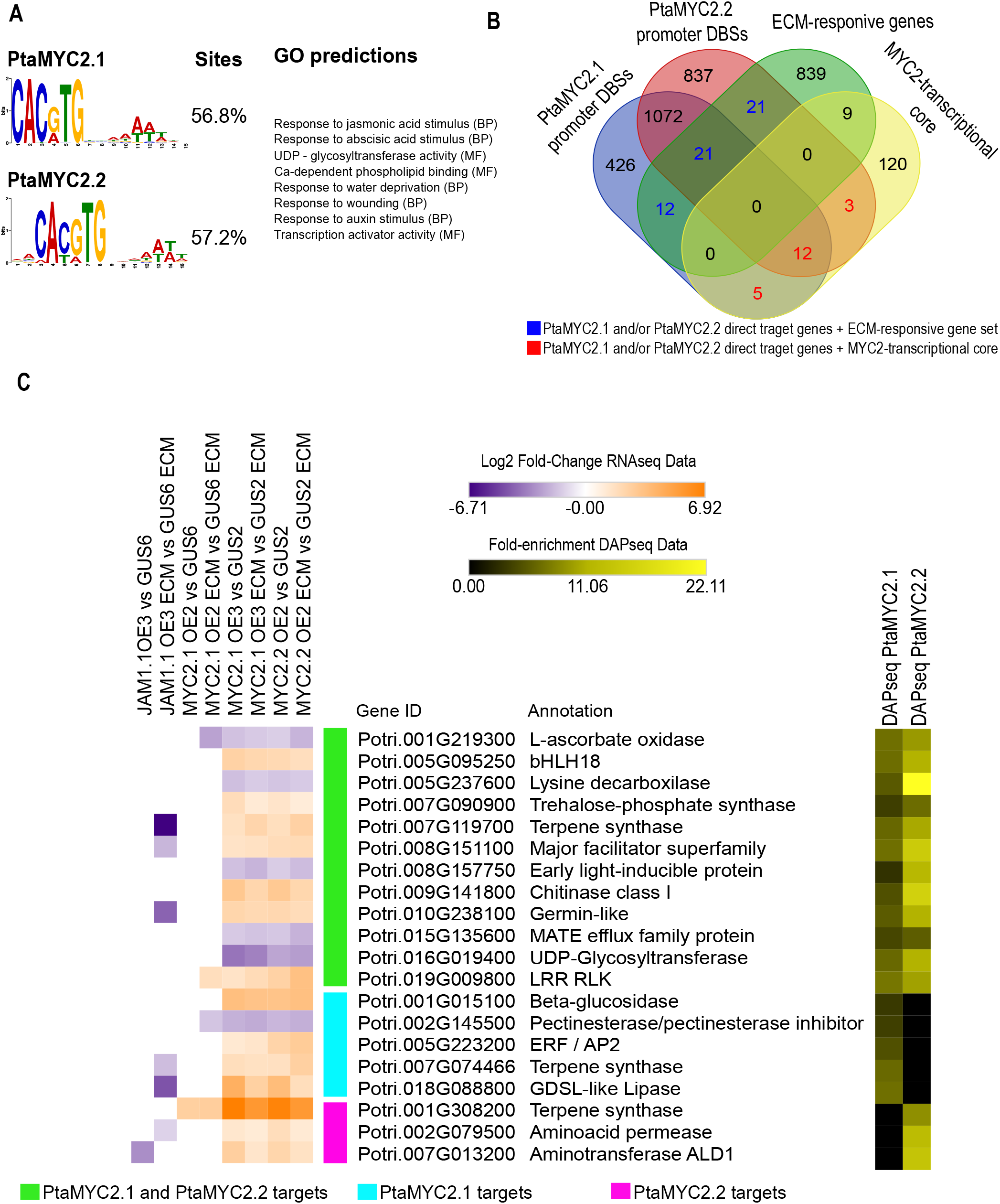
PtaMYC2.1 and PtaMYC2.2 directly target other TFs and defense-related genes important for Hartig net impairment. **(A)** Main DNA motives associated to DNA Binding Sites (DBSs) from PtaMYC2.1 and PtaMYC2.2 DAP-seq datasets, found by MEME v5.4.1. GO terms of the genes associated to the found motives were predicted by GOMo v5.4.1. **(B)** Venn’s diagram showing the overlap between the DBSs from PtaMYC2.1 and PtaMYC2.2 DAP-seq datasets and the previously RNA-seq-defined ECM-responsive genes and MYC2-transcriptional core sets. **(C)** Heatmap showing the RNA-seq differential expression (left) and DAP-seq peak fold-enrichment (right) from those genes overlapping between the DAP-seq data and the RNA-seq MYC2-transcriptional core. Manual annotation was performed according to the combination of their closest *Arabidopsis* homolog, PFAM, KEEG, KOG and GO.

Since binding events do not necessarily imply regulatory functions, we combined our results from DAP-seq and RNA-seq analyses. In order to decipher which direct MYC2 targets play a role in the HN inhibition observed in MYC2 OE lines, we overlapped PtaMYC2.1 and PtaMYC2.2 promoter DBSs with DEGs responding to fungal colonization and the MYC2-transcriptional core (Fig. 3B). Fifty-four ECM-responsive genes were directly regulated by MYC2.1 and/or MYC2.2, 49 of which were downregulated in response to ECM colonization (Fig. S11). Out of the pre-defined 149 MYC2 transcriptional core genes, 20 were directly controlled by MYC2 TFs (Fig. 3C, Dataset S21), either both (12 targets), PtaMYC2.1 only (5 targets) or PtaMYC2.2 only (3 targets). Seventeen out of 20 showed at least one G-box like motif near to the summit of their correspondent peak. When visualized, those targets exclusive of PtaMYC2.1 or PtaMYC2.2 also showed certain degree of binding events for its TF counterpart, although this was not significant (Fig. S12). Eleven out of these 20 target genes encode proteins related to biotic perception and defense functions. Three of them encoded TPSs (*PtTPS16*, Potri.007G074466 and Potri.007G119700). PtaMYC2 TFs also bound to the promoter regions of two other MYC2-transcriptional core TFs, i.e., bHLH18 (Potri.005G095250) and Apetala 2 / Ethylene Response Factor (AP2/ERF) (Potri.005G223200).

### Monoterpenes produced by poplar terpene synthases impair *in vitro L. bicolor* growth and Hartig net formation

Since TPS represented the major group of defense-related genes expressed in HN-impaired roots and several of them appeared to be directly controlled by poplar MYC2 TFs, we focused on this TPS family. There was a total of 12 differentially expressed TPS genes in at least one of our transgenic lines and/or conditions, which shared a similar pattern of expression (Fig. 4A). Moreover, we found a significant negative correlation (Pearson’s correlation, p < 0.05) between the expression pattern of regulated TPS-encoding genes (MYC2 and JAM1 OE lines vs GUS lines) and the degree of HN formation in the same transgenic lines. In summary, the more TPS-encoding genes were upregulated, the less HN developed (Fig. 4B).

**Figure 4.**
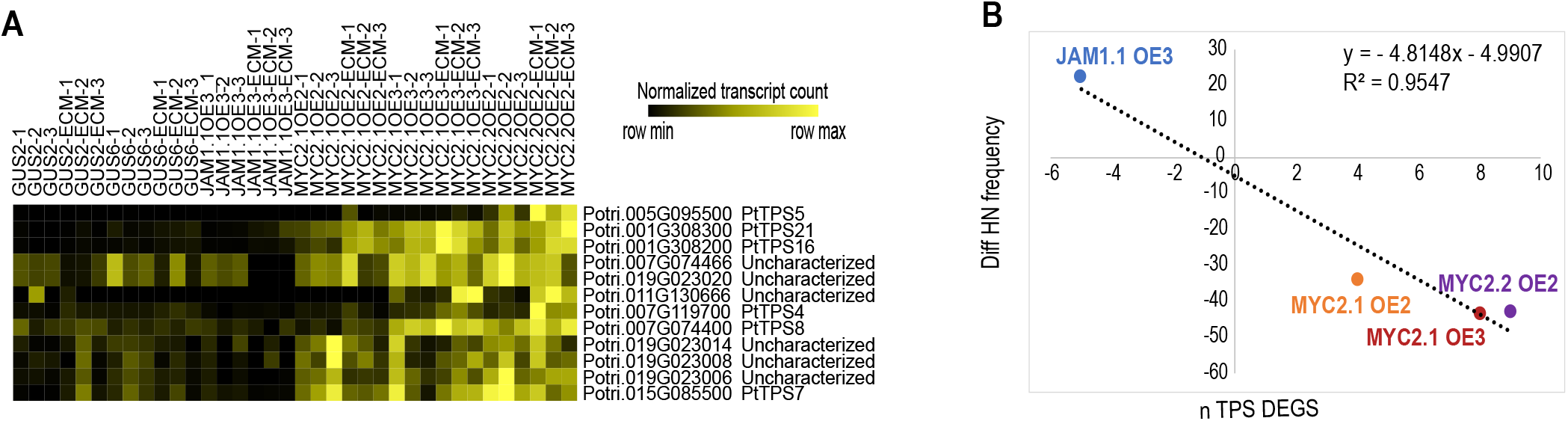
Expression patterns of root terpene synthases are correlated to Hartig net impairment. **(A)** DESeq2-normalized transcript counts for each TPS-encoding gene that was differentially expressed in at least one of the MYC2 OE lines and conditions, according to RNA-seq results. **(B)** Linear regression between the number of TPSs differentially expressed in MYC2.1 OE2, MYC2.1 OE3, MYC2.2 OE2 and JAM1.1 OE3 (x axis) and the differential of HN frequency of each of those lines with respect to the GUS controls (y axis). The trend of TPS regulation is represented by positive (upregulated) or negative (downregulated) numbers.

The known products of PtTPS16 (Potri.001G00308200) and PtTPS21 (Potri.001G00308300) activity include γ-terpinene, (-) limonene, (-) camphene, (-) α-pinene and (-) β-pinene. We thus assessed the effect of these terpenes on *L. bicolor* growth and ECM formation. A mix of these monoterpenes inhibited fungal growth *in vitro* five days post application. The inhibition rate ranged from ∼100% for 0.5 μL cm^-3^ to 20% for 0.0025 μL cm^-3^ (Fig. 5A). Individually, an inhibitory effect was observed for (-) camphene (0.1 μL cm^-3^) and, to a higher extent, for (-) α- and (-) β-pinene (0.1 μL cm^-3^). In contrast, (-) limonene or γ-terpinene (0.1 μL cm^-3^) had no detrimental effect, but led to a weak but significant increase in growth rate (Fig. 5B). Fungal growth was recovered after the total evaporation of the volatile monoterpenes (Fig. 5C). Of note, such growth inhibition was also obtained when the same monoterpene mix was applied to other ECM and endophyte fungi isolated from the poplar rhizobiome (39) (Fig. 5D, Fig. S13), although it was less severe for fungal endophytes. The application of the monoterpene mix not only inhibited fungal growth, but also affected plant phenotype and HN formation in a time-dependent manner (Fig. 6). Plant phenotype and fungal colonization (21 dpi) were assessed in the presence of the monoterpene mix (0.5 μL cm^-3^) at two different time points of the interaction: early interaction with only formation of the mantle (7 dpi) and mature interaction with presence of HN (14 dpi). A decrease in shoot fresh weight, lateral root number, ECM rate and HN formation was observed when the monoterpene mix was applied at 7 dpi, but its effect was less pronounced when applied at 14 dpi (Fig. 6). The opposite trend was observed with HN initiation points, although no significant differences were found and HN depth was not affected in plants treated at 7 dpi, but was greater for those treated at 14 dpi (Fig. 6).

**Figure 5.**
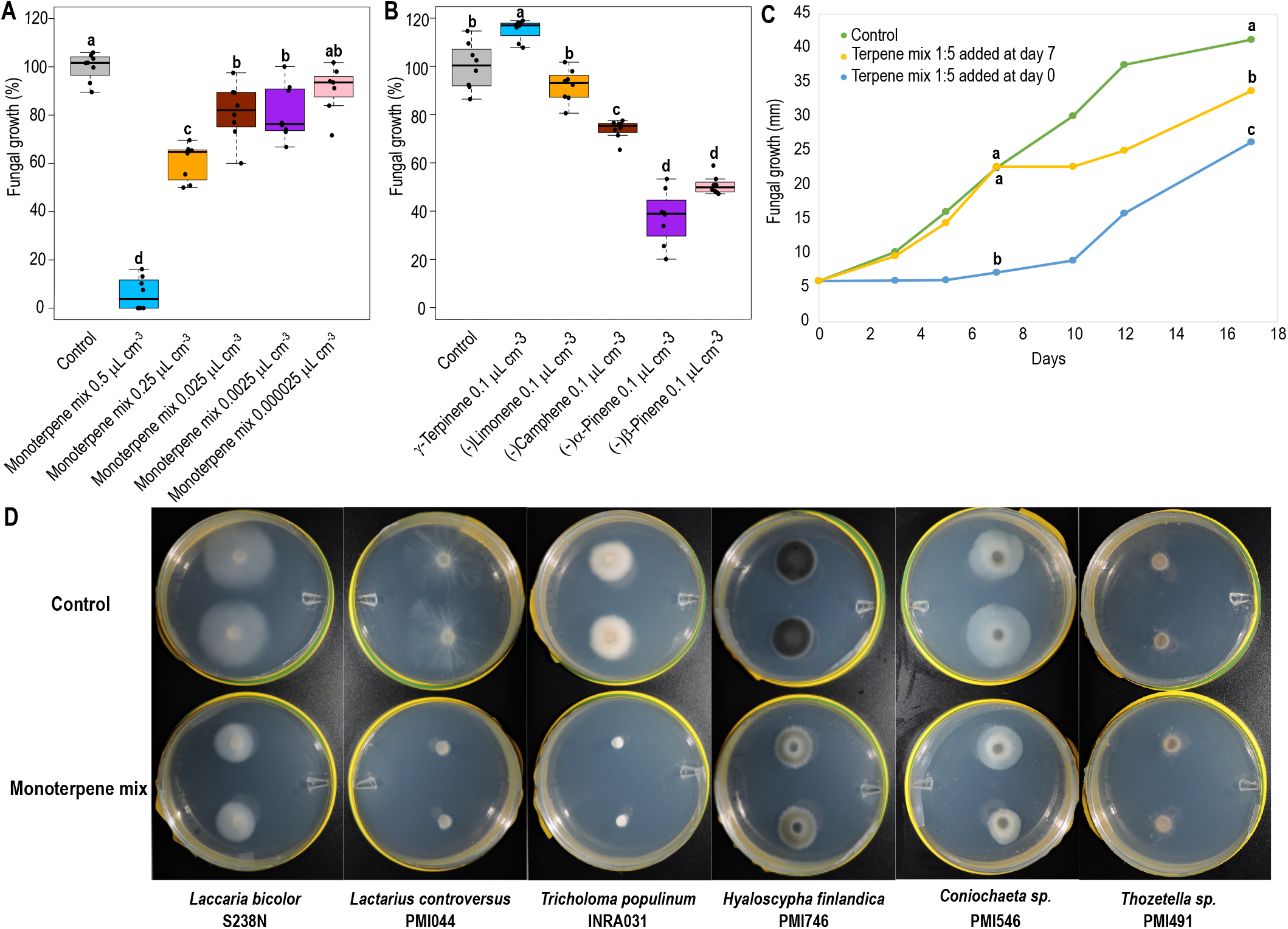
Monoterpenes produced by PtTPS16 and PtTPS21 inhibit *in vitro* fungal growth. Boxplots represent *in vitro* fungal growth normalized to the maximal growth (100%) when treated with **(A)** different concentrations of the monoterpene mix or **(B)** isolated monoterpenes. Whiskers represent the limits of the 1.5 interquartile range. Different letters represent significant differences based on Kruskal-Wallis one-way analysis of variance and Least Significant Difference (LSD) post-hoc test (p < 0.01), n=8. **(C)** Each line represents fungal growth when the monoterpene mix was applied at different time points, n=8. **(D)** Representative images of different fungi grown on agar plates containing MMN media with and without the monoterpene mix.

**Figure 6.**
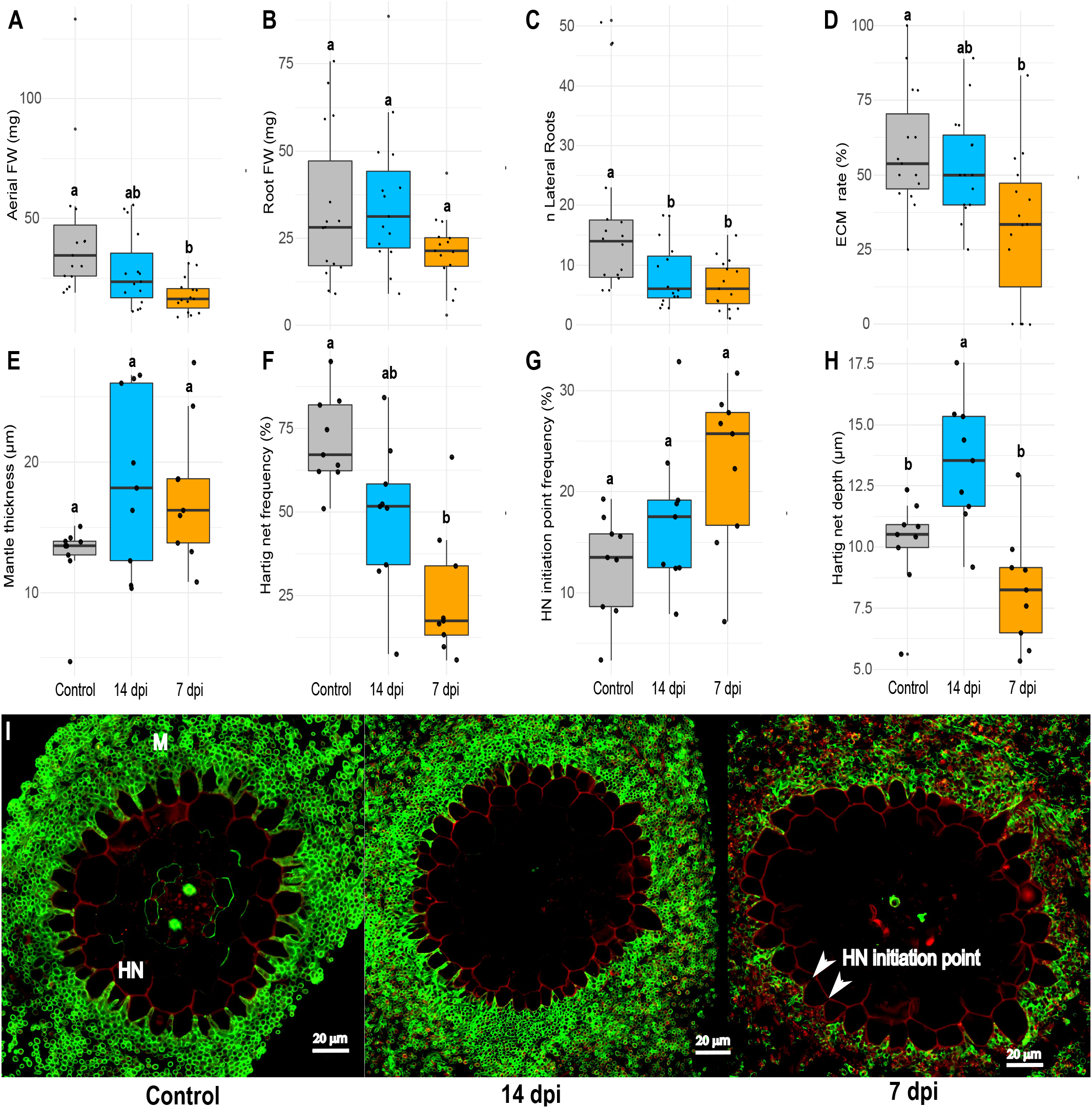
Monoterpenes produced by PtTPS16 and PtTPS21 affect plant phenotype and ECM formation in a time-dependent manner. Boxplots represent the (**A**) shoot and (**B**) root fresh weights, **(C)** number of lateral roots, **(D)** ratio between ECM lateral root tips and total lateral root tips (ECM rate), **(E)** fungal mantle thickness, **(F)** ratio between the apoplastic space surface colonized by *L. bicolor* and total apoplastic space surface (HN frequency), **(G)** Hartig net initiation points and **(H)** Hartig net depth. All measures were performed on poplar root tips inoculated with *L. bicolor* during 21 days and treated with the monoterpene mix at 7 or 14 dpi. Whiskers represent the limits of the 1.5 interquartile range. Different letters represent significant differences based on Kruskal-Wallis one-way analysis of variance and Least Significant Difference (LSD) post-hoc test (p < 0.01), n=15 (A-D), n=9 **(E-H). (I)** Representative confocal microscopy images of transversal sections of roots at 21 dpi with *L. bicolor* and treated with the monoterpene mix added at 7 or 14 dpi. White arrows indicate examples of Hartig net (HN) initiation points. Propidium iodide (red) stains the root cell wall, while AlexaFluor® WGA-288 (green) stains the fungal cell-wall. Scale bar: 20 μm.

## Discussion

The ability of MYC2 to control other TFs and multiple defensive genes has been widely reported for *Arabidopsis*, denoting a major role for MYC2 in defense against necrotrophic pathogens and herbivores (17). Our data shows that this pathway is also of relevance for the establishment of the ECM symbiosis in poplar roots, with special interest in the control of poplar MYC2s on TFs that belong to the MYC2-transcriptional core, i.e., *ERF/AP2* (Potri.005G223200) and *bHLH18* (Potri.005G095250). The direct control of *ERF/AP2* expression by MYC2 is likely a mechanism allowing the cross-talk between JA and ET signaling pathways in ECM symbiosis (40), while the control of bHLH18, which has similar binding affinities to those of PtaMYC2.1 and PtaMYC2.2, suggests an additive role for this TF, similar to the interplay between MYC2 and MYC3 in *Arabidopsis* (41). Poplar MYC2s also directly control the expression of several defense-related genes during *Populus x Laccaria* ECM development. These genes are well known to be involved in plant defense reactions in other plant species, such as LRR-RLKs, ALD1, germin-like proteins, chitinases, β-glucosidases, GDSL-like lipases, trehalose-phosphate synthase and TPSs (42-49). Notably, the degree of ECM inhibition appeared to be quantitatively correlated to the relative expression of these defense-related genes, as exemplified by the negative correlation between the number of upregulated TPS-encoding genes and the inhibition of HN formation. Surprisingly, a TPS encoding gene, *PtTPS21*, is also slightly upregulated during ECM development, but its expression was more than 10 times higher in MYC2 OE lines. This suggests that only a weak and tight upregulation of plant defenses is required in ECM root tips, likely to impede fungal over-colonization of the roots, but an excessive upregulation in MYC2 OE lines resulted in impairment of HN formation. The expression of defense genes such as *PtTPS21, PtTPS16* and other TPS, chitinases and endopeptidase inhibitors was strongly upregulated in the presence of Me-JA and at one week post *L. bicolor* inoculation, but repressed two weeks post-inoculation (40). These findings suggest that both the extent and timing of this defense reaction repression are critical for the final outcome of ECM establishment.

Unlike *Arabidopsis, Populus* possess two different homologs of MYC2. Our results indicate that the activity of both these orthologs should be turned off to allow HN establishment, since the overexpression of either one impaired the full HN formation. These genes are both expressed in roots and they shared a high proportion of their promoter DBS. As reported for *Arabidopsis* MYC2, MYC3 and MYC4, we speculate that PtaMYC2.1 and PtaMYC2.2 have overlapping function in some signaling cascades, such as those involved in ECM establishment, but can also play distinct roles in other processes or organs (50-51). Additionally, PtaJAM1.1 might counteract the activities of PtaMYC2.1 and PtaMYC2.2, as observed in *Arabidopsis* (32). The sequence similarity of the PtaJAM1.1, PtaMYC2.1 and PtaMYC2.2-DNA binding domain (37) and the fact that JAM1.1 OE3-downregulated genes were enriched in the same defensive functions than those of the MYC2-transcriptional core upregulated genes suggest that this antagonistic effect is the result of PtaJAM1.1 direct binding to PtaMYC2s DBSs.

In summary, here we suggest a working model that extents our previously proposed hypothesis (34) (Fig. 7). In poplar ECM root tips, the effector LbMiSSP7 enters root cells and migrates to the nuclei where it binds to and stabilizes the JA co-receptor PtaJAZ6, counteracting PtaMYC2s activity (35-37). Using transgenic poplars, we showed that both PtaMYC2 homologs coordinate the root cell transcriptional reprogramming that would hinder *in planta* fungal growth, if not controlled by the MiSSP7-JAZ6 complex. This process mainly takes place through the control of the upregulation of other TFs and specific defense-related genes. Among these defense-related genes, we showed that root TPSs play a key role, as the volatile compounds produced by these MYC2-regulated TPSs inhibit ECM formation as well as the growth of *L. bicolor* and other ECM and endophytic fungi isolated from the poplar rhizosphere. The roles of volatile terpenes in mediating above-ground interactions between plants and other living organisms are very well documented (52, 53). By contrast, little is known about the role(s) of terpenes produced by roots on the belowground microbial communities (54). The products of PtTPS16 an PtTPS21, in addition to the product of the sesquiterpene synthase PtTPS5, have previously shown inhibitory activity against oomycete ingress in poplar roots (37, 55). Other previously characterized poplar TPSs play a role in leaf herbivore defense (56, 57), although our results suggest that they could also be involved in root defense against fungi. A recent study highlighted the role of *Arabidopsis* root triterpenes in the selection, assembly and maintenance of the root bacterial microbiota (58). Our study demonstrates the role of MYC2 in ECM symbiosis and at the same time provides the first evidence of a role of poplar root monoterpenes in modulating a belowground mutualistic interaction. Of note, it has been previously shown that fungal sesquiterpenes act as signaling molecules during ECM formation (59). Overall, those data support the contention that terpenes are used as a chemical language between roots and soil-borne ECM fungi. Future studies will aim to follow the spatiotemporal pattern of MYC2-coordinated defenses, with especial focus on TPSs, during fungal ingress to better understand their role during this developmental process, as we showed that the fungal colonization depends on the maintenance of their repression.

**Figure 7.**
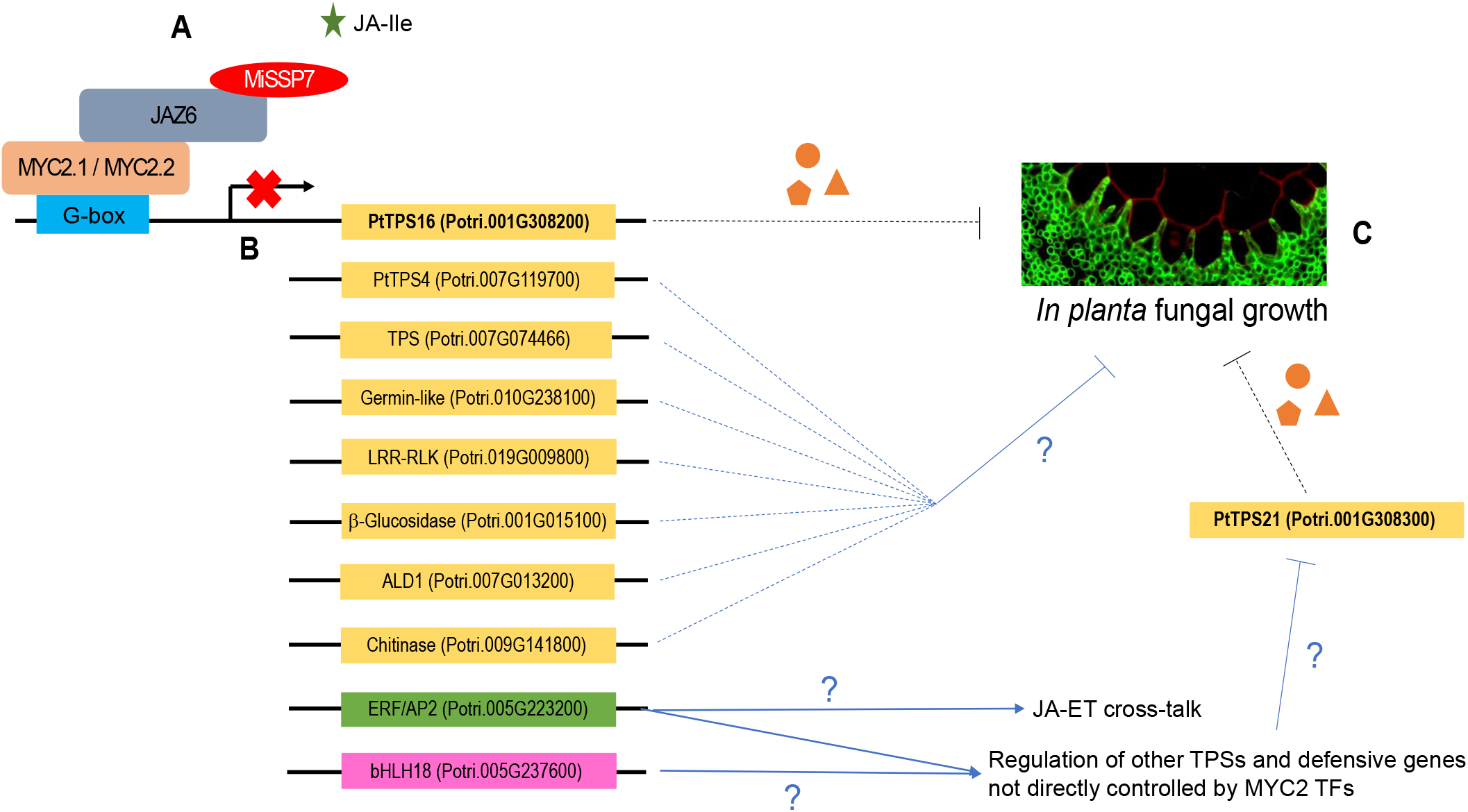
Proposed working model of action for *Populus* MYC2 transcription factors during ectomycorrhiza (ECM) development. **(A)** In *Populus* ECM root tips, *L. bicolor* MiSSP7 enters root cells and migrates to nuclei where the symbiotic effector binds to the JA co-receptor PtaJAZ6, preventing its degradation upon JA-Ile perception (34-35). MiSSP7 not only prevents the degradation of PtaJAZ6, but also strengthens the interaction between PtaJAZ6 and poplar PtaMYC2s (37). **(B)** PtaMYC2.1 and PtaMYC2.2 directly control the expression of several poplar defense-related genes encoding TPS, chitinase I, germin-like, LRR-RLK, β-glucosidase or ALD1 and other TFs, such as ERF/AP2 or bHLH18. The interaction between PtaJAZ6 and both PtaMYC2.1 and MYC2.2 TFs allows the development of the ECM symbiosis by avoiding the activation of these genes, that would otherwise impair *in planta* fungal growth. **(C)** MYC2-regulated TPS volatile products can inhibit fungal growth and ECM formation *in vitro*.

## Materials and methods

### Generation of poplar stable transgenic lines

We isolated the *P. tremula x alba* full-length coding sequence (CDS) of *PtaMYC2*.*1* (Potri.001G142200), *PtaMYC2*.*2* (Potri.003G092200) and *PtaJAM1*.*1* (Potri.014G099700) by PCR, using as template a cDNA library obtained from poplar-*L*.*bicolor* ECM, plus a fragment of the *GUS* gene (bases 345-1034) used as negative control. We cloned these sequences into the pJCV53 overexpression vector (VIB-Plant Systems biology; Ghent University) using the Gateway® cloning system (Thermo Fisher Scientific) and generated *Populus x tremula alba* 717-1B4 transgenic stable lines by the agrotransformation method according to Cope et al. (60). The list of primers used for cloning for this and other purposes is provided in Dataset S22 and further details of the protocol are provided in SI Appendix, Materials and Methods.

### Generation of poplar composite plants and localization of PtaMYC2.1 and PtaMYC2.2 expression

We synthetized ∼2 kb before TSS of both PtaMYC2.1 and PtaMYC2.2 (GeneCust) and cloned them into the pKGWFS7 transcriptional reporter vector (VIB-Plant Systems biology; Ghent University), plus the empty vector as a control, using the Gateway® cloning system (Thermo Fisher Scientific), with its 3’-end fused to the *EGFP* and *GUS* reporter genes. Composite poplar plants with transformed roots were prepared according to Neb et al. (61), with some modifications. After 21 days of growth, roots were observed under Discovery V.8 stereomicroscope (Zeiss) coupled to a HXP120v fluorescence lighting device (Zeiss). Roots were observed under bright field and with green filter (470 nm excitation, 525 nm emission) and the number of adventitious and lateral roots that presented green fluorescence was annotated. Further details of the protocol are provided in SI Appendix, Materials and Methods.

### Plant and ECM phenotype of *P. tremula x alba* transgenic lines

Two separate experiments were carried out including GUS6, MYC2.1 OE2 and JAM1.1 OE3 (Experiment 1) or GUS2, MYC2.1 OE3 and MYC2.2 OE2 (Experiment 2). Plant and fungal preparation, and analysis of mycorrhizal phenotypes were made according to Basso et al. (40), with minor modifications. Further details of this protocol are provided in SI Appendix, Materials and Methods.

### RNA extraction and RNA-seq

Coinciding with ECM transgenic line phenotype experiments, RNA extraction from colonized and uncolonized root systems was performed according to Basso et al. (40). From these RNA samples, libraries were generated and sequenced by the Joint Genome institute using Illumina’s TruSeq Stranded mRNA HT sample prep kit and Illumina NovaSeq 6000. A detailed protocol of RNA extraction and RNA-seq downstream bioinformatic analyses is available in SI Appendix, Materials and Methods.

### DAP-seq

Open reading frames for the transcription factors PtaMYC2.1, PtaMYC2.2 and PtaJAM1.1 were cloned into pIXHALO (ABRC, Ohio State University) and expressed *in vitro* using TnT® SP6 High-Yield Wheat Germ Protein Expression System (Promega). We used empty vector pIXHALO as control. For DAP-seq experiments and analysis we followed the protocol described by Bartlett et al. (40), with minor modifications. A more detailed protocol of this process and downstream bioinformatic analyses can be consulted in SI Appendix, Materials and Methods.

### Monoterpene bioactivity assays

The monoterpenes γ-terpninene, (-)limonene, (-)camphene, (-)α-pinene and (-)β-pinene (Sigma-Aldrich, US) were exogenously applied to *L. bicolor* S238N, *Lactarius controversus* PMI044, *Tricholoma populinum* INRA031, *Hyaloscepha finlandica* PMI746, *Coniochaeta* sp. PMI546 and *Thozetella* sp. PMI491 in pure *in vitro* cultures. A terpene mix was prepared by mixing 10 μL of each pure compound (> 98%) for a total of 50 μL. Terpene mix was added in sterilized open PCR tubes included in 100 cm^3^ plates making a final terpene concentration of 0.5 μL cm^-3^, unless otherwise stated in the text. Plates were sealed with a double layer of parafilm and tape. Fungal growth of 8 independent colonies was measured after 5 days. For the quantification assay, terpene mix was submitted to sequential 1:2 and 1:10 dilutions in mineral oil, maintaining 50 μL of final volume. For the individual monoterpene assay 10 μL of each compound was mixed with 40 μL of mineral oil. For the kinetic study, fungal growth was monitored at days 3, 5, 7, 10, 12 and 17. The same procedure was followed to test their effect in the interaction between *P. tremula x alba* 717-1B4 and *L. bicolor*, maintaining the concentration of 0.5 μL cm^-3^ (150 μL of final volume in three PCR tubes). Plant and fungal preparation and analysis of mycorrhizal phenotypes were performed as for the transgenic lines.

## Acknowledgments

This research work was sponsored by the Plant- Microbe Interfaces Scientific Focus Area (http://pmi.ornl.gov) in the Genomic Science Program, the Office of Biological and Environmental Research in the U.S. Department of Energy Office of Science (AK, CVF, FM, JEMG, RV, VB), the Laboratory of Excellence ARBRE (ANR-11-LABX-0002-01) (CVF, VB, JEMG). Oak Ridge National Laboratory is managed by UT-Battelle for the U.S. DOE Office of Science (DE-AC05-589 00OR22725). The work (proposal: 10.46936/10.25585/60001022) conducted by the U.S. Department of Energy Joint Genome Institute (https://ror.org/04xm1d337), a DOE Office of Science User Facility, is supported by the Office of Science of the U.S. Department of Energy operated under Contract No. DE-AC02-05CH11231. RNA seq data and analysis were performed by KB, KK, JJ, VS and IG. Sequencing from the DAP-seq experiment was performed by GenomEast platform hosted at the IGBMC and we acknowledge Bernad Jost and Stéphanie Le Gras for their help.

