## Supplementary materials and methods for "The establishment of *Populus* x *Laccaria bicolor* ectomycorrhiza requires the inactivation of MYC2 coordinated defense response with a key role for root terpene synthases"

**SI appendix: Materials and methods**

**Generation of poplar transgenic lines**

After plasmid construction, we generated poplar transgenic lines according to Cope et al. (1). Briefly, *Rhizobium rhizogenes* (formerly *Agrobacterium rhizogenes*) Arqua 4 competent cells were transformed by electroporation with OE vectors. After selection of transformed colonies, *R. rhizogenes* strains were cultivated on 5 mL of liquid MG/L broth (5 $g.L^{-1}$ tryptone, 2.5 $g.L^{-1}$ yeast extract, 5.2 $g.L^{-1}$ NaCl, 2 $g.L^{-1}$ glutamic acid, 10 $g.L^{-1}$ mannitol, 0.2 $g.L^{-1}$ $MgSO_4$-$7H_2O$, 0.5 $g.L^{-1}$ $K_2HPO_4$, pH 7.0, with addition of 4 $\mu g.L^{-1}$ biotin, 50 $\mu g.mL^{-1}$ rifampicin and 50 $\mu g.mL^{-1}$ spectinomycin after autoclaving) in shaker overnight (O/N) at 28°C and 200 rpm. The following day, the culture was brought up to 50 mL by addition of fresh MG/L broth and grown in shaker O/N at 28°C and 200 rpm. Cultures were centrifuged at 3500 rpm for 20 minutes and the bacterial pellet was resuspended in 50 mL of induction broth (modified Murashige & Skoog with 1X micronutrients (Sigma-Aldrich), 1X macronutrients (Sigma-Aldrich), 0.25 $g.L^{-1}$ MES (Sigma-Aldrich), 0.1 $g.L^{-1}$ myo-inositol, 30 $g.L^{-1}$ sucrose, pH 5.8, with addition of 100 mM acetosyringone (Sigma-Aldrich) after autoclaving). Bacterial cultures were diluted in induction broth to an $OD_{600}$ of 0.8. Leaves from *in vitro* poplar plants were excised with scissors and incisions in prominent veins on the underside of the leaf were practiced with a scalpel. Wounded leaves were kept in sterile water to prevent desiccation. Leaves were removed from the water, immersed in the bacterial culture for 5 minutes and placed on co-cultivation solid medium (modified Murashige & Skoog previously described with 7$g.L^{-1}$ agar agar) and incubated in the dark at 24°C for 2 days. Afterwards, leaves were transferred to antibiotics medium (co-cultivation medium, without acetosyringone but with cefotaxime and carbenicillin at 250 $\mu g.mL^{-1}$ each (Duchefa Biochemie)) to eliminate excess bacteria. Leaves were kept at 24°C, 60% humidity, 50 $\mu mol.m^{-2}.s^{-1}$ light intensity under a 16-h photoperiod and transferred onto new antibiotics medium every 2 weeks until emergence of roots from the veins of the leaves. Leaves were then placed on selection medium (antibiotics medium, with addition of 50 $\mu g.mL^{-1}$ kanamycin (Sigma-Aldrich)) until the roots reached at least 2-3 cm of length. Roots were observed under a fluorescent stereomicroscope to detect the red fluorescence of the mRFP or dsRED transformation markers. Transformed roots were excised from the leaves and placed on regeneration medium (selection medium, with addition of 0.055 $\mu g.mL^{-1}$ thidiazuron (Sigma-Aldrich)) and cultivated in the same conditions until emergence of shoots. Shoots derived from the same transformed root were considered as belonging to the same transgenic line and further propagated in the same conditions as wild-type poplar plants.

**Generation of poplar composite plants with transformed roots**

We generated composite poplar plants with transformed roots according to Neb et al. (2), with some modifications. Briefly, *R. rhizogenes* Arqua 4 competent cells were transformed by electroporation with transcriptional reporter plasmids. After selection of transformed colonies, *R. rhizogenes* strains were cultivated on liquid YEB (5g.$L^{-1}$ beef extract, 1 g.$L^{-1}$ yeast extract, 5 g.$L^{-1}$ peptone, 5 g.$L^{-1}$ sucrose, 0.5 g.$L^{-1}$ $MgCl^2$ , pH 7.2, add 50 µg.$mL^{-1}$ of rifampicin (Sigma-Aldrich) and 50 µg.$mL^{-1}$ of spectinomycin (Sigma-Aldrich) after autoclaving) O/N at 28°C and 200 rpm. Afterwards, 250 µL of culture was plated on solid YEB plates and kept in static oven at 28°C for 3 days. The thick layer of cells was collected with the help of a scalpel and suspended in 3 mL of sterile water in order to generate foam. Axenically grown wild-type poplar cuttings were briefly dipped in the bacterial foam and transferred to co-cultivation plates (modified Murashige & Skoog with 1X micronutrients (Sigma-Aldrich), 0.5X macronutrients (Sigma-Aldrich), 3% sucrose, 1 g.$L^{-1}$ MES (Sigma-Aldrich), 20 mg.$L^{-1}$ L-glutamine, 0.1 mg.$L^{-1}$ Ca-panthotenate, 0.1 mg.$L^{-1}$ L-cysteine, 0.01 mg.$L^{-1}$ biotine, and 1X Gamborgs vitamins, pH 5.8). Plates were sealed and kept in the dark at 24°C with 60% humidity for 3 days. Subsequently, plants were transferred onto co-cultivation plates with addition of Amoxycillin sodium / Clavulanate potassium at 250 µg.$mL^{-1}$ each (Duchefa Biochemie) in order to eliminate excess growth of *R. rhizogenes*. Plants were kept at 24°C with 60% humidity and 50 µmol.$m^{-2}$.$s^{-1}$ light intensity under a 16-h photoperiod. Plants were transferred onto new co-cultivation plates with antibiotics until the emergence of roots.

**Plant and fungal materials, growth conditions and poplar- *L. bicolor* co-cultivation method**

Plant and fungal materials were cultured as described by Felten et al. (3). Briefly, transgenic grey poplar (*Populus tremula x Populus alba* line INRA 717-1-B4) clones were micropropagated *in vitro* and grown in half-strength Murashige and Skoog (MS 1/2) medium in glass culture tubes in a growth chamber at 24°C and 150 µmol.$m^{-2}$.$s^{-1}$ light intensity under a 16-h photoperiod. Light came from OSRAM Fluorescent tubes (50/50 Fluora / Cool white) placed 15 cm from poplar plants. The dikaryotic vegetative mycelia of strain S238N of the ectomycorrhizal fungus *Laccaria bicolor* were maintained on modified Pachlewski agar medium P5 at 25°C in the dark (4). For *in vitro* co-culture of poplar with *L. bicolor*, we used the sandwich system described by Felten et al. (3). Briefly, ~10-mm-long rooted stem cuttings from *in vitro*-grown poplar plants were transferred to Petri dishes containing Pachlewski agar medium with reduced sugar (P20) covered with a cellophane membrane and a second, mycelium-covered cellophane membrane was placed on the roots (colonized plants). For single cultures, poplar plants were grown in absence of fungal mycelium (uncolonized plants). The Pachlewski agar medium was supplemented with MES-Na to maintain the pH at 5.8. The Petri dishes were positioned vertically and incubated for 3 weeks in a growth chamber at 20°C and 50 $\mu mol.m^{-2}.s^{-1}$ light intensity under a 16-h photoperiod.

**Analysis of mycorrhizal phenotypes**

The quantification of ectomycorrhizal colonization and the observation of ectomycorrhizal structures were performed as described by Felten et al. (3). Briefly, colonized plants were observed under a Discovery V.8 stereomicroscope (Zeiss), and short, rounded lateral roots surrounded by fungal mantle were considered to be colonized. The rate of ectomycorrhizal colonization was defined as the ratio of colonized lateral roots to the total number of lateral roots (expressed as a percentage). Between 18 and 24 plants per transgenic line were observed. To confirm the development of intraradical Hartig net, at least six ECM per transgenic line were fixed with 4% paraformaldehyde in 1X phosphate-buffered saline and embedded in 4% (w/v) agarose. Transverse 30-µm-thick sections of ECM obtained 200 to 400 µm from the lateral root tip were produced with a Leica VT1200 S vibratome and stained with propidium iodide (1:100 dilution; Sigma-Aldrich) and WGA-488 (1:100 dilution; Wheat Germ Agglutinin, Alexa Fluor™ 488 Conjugate, Thermo Fischer Scientific) to highlight plant and fungal structures, respectively. The sections were observed under a Zeiss LSM 700 laser scanning microscope and the images analyzed with Fiji software (5). Root diameter, fungal mantle thickness, and Hartig net depth were measured. Hartig net frequency (expressed as the percentage of root apoplastic spaces occupied by fungal hyphae) and frequency of Hartig net initiation (expressed as the percentage of root apoplastic spaces starting to be occupied by fungal hyphae) were also calculated. At least three sections per sample were analyzed.

**RNA extraction and RT-qPCR**

For the transgenic poplar lines, the whole root systems from three colonized or uncolonized poplar plants were pooled, snap-frozen in liquid nitrogen and stored at -80°C for subsequent RNA extraction. RNA was extracted with the plant RNeasy Mini Kit (Qiagen), following manufacturer's instructions. RNA quality was assessed with a Fragment Analyzer Automated CE System (Agilent). Five hundred nanograms of RNA were used to synthesize cDNA with the iSCRIPT™ cDNA Synthesis Kit (BioRad) and the resulting cDNA was diluted to 2.5 ng/mL in RNase-free water. Two microliters of diluted cDNA were added to 5 µL of iQ SYBR Green Supermix (BioRad) and 0.5 µL of 10 nM forward and reverse primers in a 10 µL reaction. Real-time qPCR was performed in a Rotor-Gene Q thermocycler (Qiagen) with two technical replicates. Threshold cycles were calculated for each pair of tested primers and the number of cDNA molecules for the gene of interest present at the beginning of the reaction was estimated with the following formula:

$$Numberofmolecules = E_p^{-Ct}$$

with $E_p$ = coefficient of primer pair efficiency (equals 2 in case of 100 % efficiency) and Ct = threshold cycle. The expression level of each gene of interest was normalized by the expression level of the housekeeping gene Ubiquitin (*UbiQ*) for each sample. For the screening of transgenic poplar lines, single shoot replicates have been used. To test colonized and uncolonized poplar roots belonging to the transgenic lines, two or three biological replicates have been produced. The list of primers used for RT-qPCR is provided in Dataset S22.

**RNA-seq and differential expression analysis**

RNA was extracted as detailed above. From these RNA samples, libraries were generated and sequenced by the Joint Genome institute using Illumina's TruSeq Stranded mRNA HT sample prep kit and Illumina NovaSeq 6000. Plate-based RNA sample prep was performed on the PerkinElmer Sciclone NGS robotic liquid handling system using Illumina's TruSeq Stranded mRNA HT sample prep kit utilizing poly-A selection of mRNA following the protocol outlined by Illumina in their user guide (https://support.illumina.com/sequencing/sequencing_kits/truseq-stranded-mrna.html), and with the following conditions: total RNA starting material was 100 ng per sample and 10 cycles of PCR was used for library amplification. The prepared libraries were then quantified using KAPA Illumina library quantification kit (Roche) and run on a LightCycler

480 real-time PCR instrument (Roche). The quantified libraries were then multiplexed and the pool of libraries was then prepared for sequencing on the Illumina NovaSeq 6000 sequencing platform using NovaSeq XP v1 reagent kits (Illumina), S4 flow cell, following a 2x150 indexed run recipe. Raw reads were analyzed using FASTQC for quality control (Andrews et al., 2010), paired, trimmed, and mapped against *Populus trichocarpa* v4.1 transcripts (https://phytozome-next.jgi.doe.gov/info/Ptrichocarpa_v4_1) using CLC Genomics WORKBENCH20 (Qiagen,Courtaboeuf, France). For mapping, the minimum length fraction was 0.9, the minimum similarity fraction 0.9, and the maximum number of hits for a read was set to 10. Before performing further analyses, reads that mapped in genes that contained in average less than 5 raw mapped reads across all the conditions were removed. Raw read counts were normalized (with the function *counts* in DESeq2), and differential transcription levels of the contigs (FDR adjusted $p < 0.05$) were calculated with DESeq2 (6). The RNA-seq data were consistent according to the quality assessment (Fig. S14), permitting our transcriptomic analyses. Condition-specific differentially expressed genes (log2(fold-change) > 1 or < −1, false discovery rate FDR < 0.05) were identified using a custom R script. To identify ECM responsive genes, we compared each separate line in colonized vs uncolonized conditions. To identify genes related to the overexpression, we compared each OE lines with a GUS control in both colonized and uncolonized conditions. Since samples clustered together according to the experimental setup, we decided to use GUS6 as control for experiment 1 and GUS2 for experiment 2 (Fig. S14).

**DAP-seq**

DAP-seq was performed following the protocol of Bartlett et al. (7), with minor modifications. Genomic DNA was extracted from *P. tremula x alba* 717-1B4 roots using DNeasy plant mini kit (Qiagen), following's manufacturer's instructions. gDNA quality was checked in 0.75% agarose gel and it was diluted to a concentration of 40 ng/µL in 128 µL, measured by Qubit dsDNA HS Assay kit (Thermofisher Scientific). gDNA was sonicated in Covaris M220 (Covaris) ultrasonicator (peak incident power: 50w; Duty factor: 20%; Cpb: 200; Time 160s). The quality of sonicated gDNA was checked in TapeStation 2200 (Agilent Technologies), confirming that the majority of the gDNA had a size between 200 and 600 bp. Sonicated gDNA was cleaned up and submitted to end repair, A-tail reaction, adaptor ligation and a final cleanup following the protocol. Verification of adaptor ligation by qPCR was performed, using an aliquot of

DNA(+adaptors) and another of DNA(-adaptors), obtaining amplification only for DNA that contained the adaptors. To study all the potential binding sites of PtaMYC2.1 and PtaMYC2.2, we removed methylation by performing PCR (11 cycles), as suggested in the protocol. Open reading frames for transcription factors PtaMYC2.1, PtaMYC2.2 and PtaJAM1.1 were cloned into pIXHALO (ABRC, Ohio State University) and expressed *in vitro* using TnT® SP6 High-Yield Wheat Germ Protein Expression System (Promega). We also included pIXHALO empty vector as a control. Halo-fused proteins were bound to Magne HaloTag beads. The success of protein in vitro expression and Magne HaloTag beads bound was measured by performing ECL Western Blot using anti-HaloTag monoclonal antibodies before and after bead binding, as described in the protocol (Fig. S9). After several washes with PBS+NP40, DNA was added, washed, and recovered following the protocol instructions. Recovered DNA that bound to the proteins of interest was amplified by PCR, pooled and loaded to a 1.5% agarose gel. Using a scalpel blade, we cut ~200 to ~600 bp DNA smear from the gel, extracted the DNA using Qiagen Gel Extraction Kit (Qiagen) and measured concentration in Qubit dsDNA HS Assay (Thermofisher Scientific). All the adaptors and primers used in this protocol can be consulted in Dataset S22.

Purified gDNA bound to PtaMYC2.1 and PtaMYC2.2 was sequenced by GenomEast platform (Strasbourg, France) using Illumina Hiseq4000, paired-end (2x100b) technology. Raw reads were submitted to quality control using FASTQC (8), Illumina primer sequences were trimmed using TRIMMOMATIC (9), resulting reads were mapped against *P. trichocarpa* v4.0 genome using BOWTIE2 (10) and filtered to obtain only unique mapped reads. MACS2 (11) was used to perform peak calling, using the empty vector as a reference. We used ChipQC R package to perform quality control over our data and then peaks were annotated using ChIP seeker and GenomicFeatures R packages (12, 13). We filtered those peaks that were present in the promotor regions of genes (<3 kb of the Transcription Starting Site) and performed motif enrichment using MEME suite v4.5.1 (14) and assigned functions to the enriched motifs using GOMo, with plant database. DAP-seq peaks were visualized using IGV viewer.

(8) S. Andrews, FastQC:  A Quality Control Tool for High Throughput Sequence Data.  [Online]. Available online at: http://www.bioinformatics.babraham.ac.uk/projects/fastqc/
