## Supplementary material for "The establishment of *Populus* x *Laccaria bicolor* ectomycorrhiza requires the inactivation of MYC2 coordinated defense response with a key role for root terpene synthases": Suplementary figures

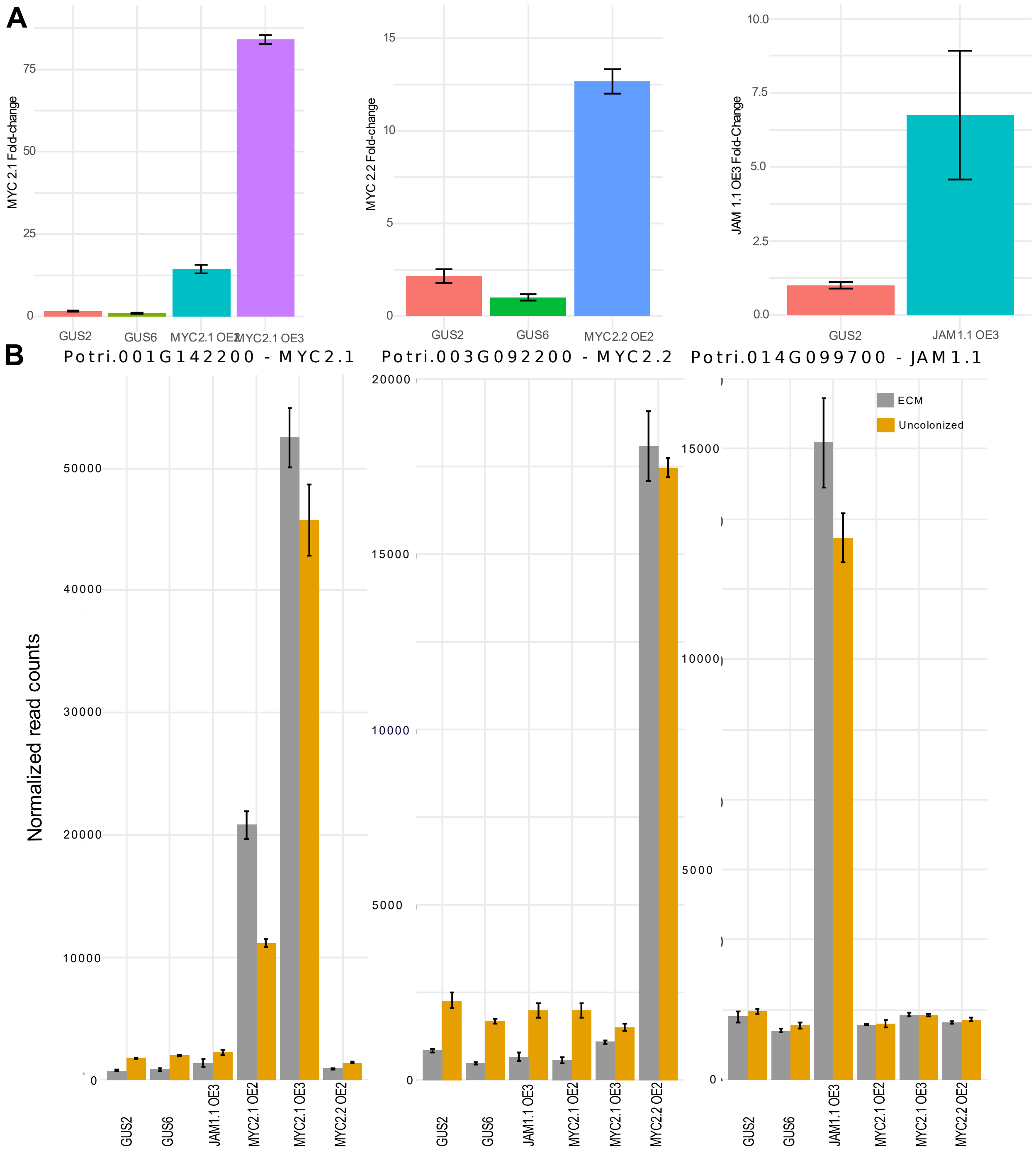


**Figure S1. Gene expression levels of transgenic lines overexpressing *PtaMYC2.1*, *PtaMYC2.2* and *PtaJAM1.1* (A)** Barplots representing the relative *PtaMYC2.1*, *PtaMYC2.2* and *PtJAM1.1* expression, as detected by RT-qPCR in shoots of control (GUS), MYC2.1 OE2, MYC2.1 OE3, MYC2.2 OE2 and JAM1.1 OE3 transgenic lines, n=3. Gene expression was normalized by the expression level of the poplar housekeeping gene Ubiquitin. **(B)** Barplots show normalized reads from RNA-seq data of PtaMYC2.1 (left), PtaMYC2.2 (center) and PtaJAM1.1 (right) in MYC2.1 OE2, MYC2.1 OE3, MYC2.2 OE2 and JAM1.1 OE3 lines in colonized and uncolonized conditions, n=3.

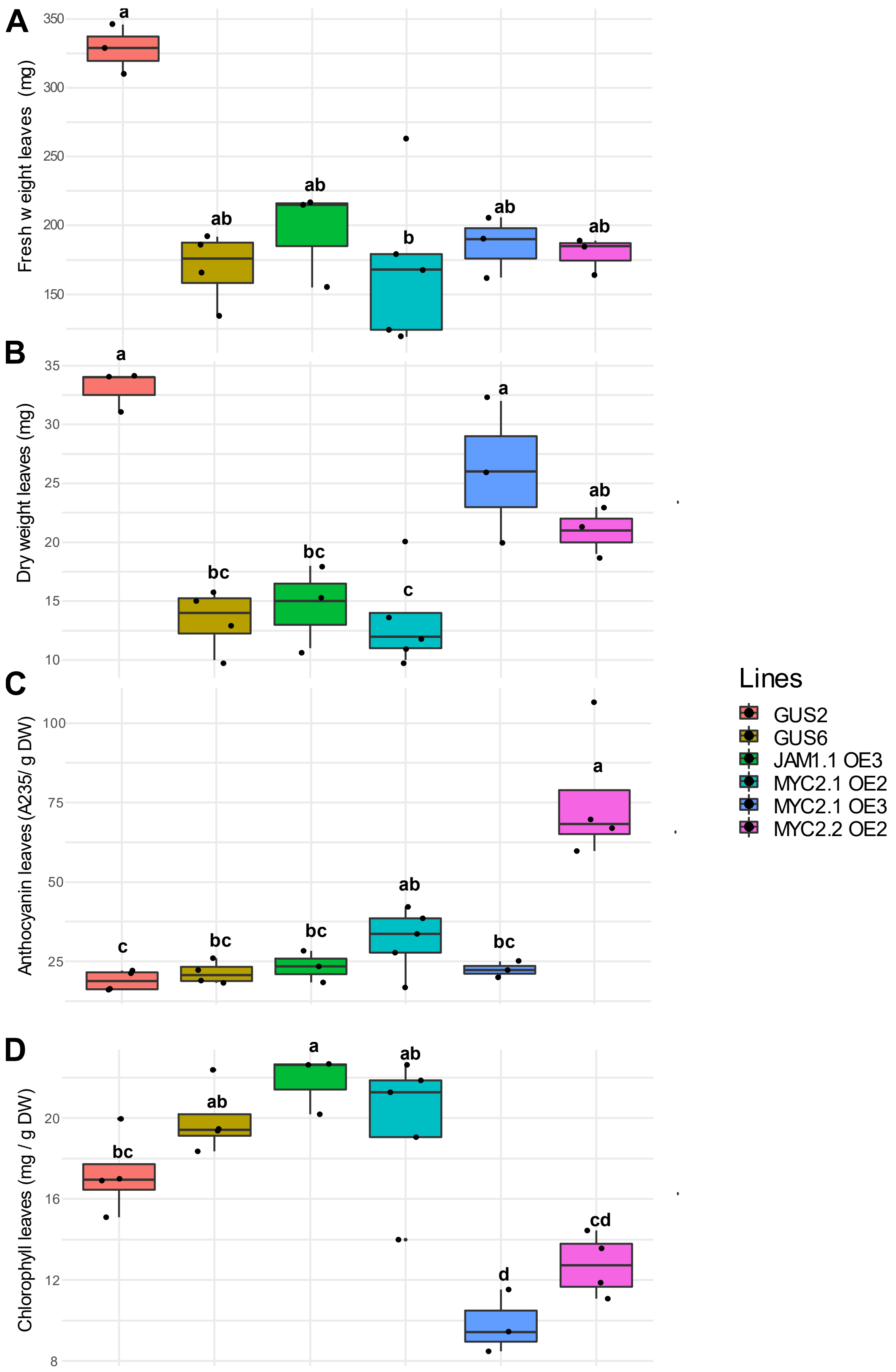


**Figure S2. The overexpression of *PtaMYC2.1*, *PtaMYC2.2* and *PtaJAM1.1* differentially alters shoot physiology.** Boxplots showing the distribution of leaf **(A)** fresh weight (FW), **(B)** dry weight, (**C**) anthocyanin and **(D)** chlorophyll content of transgenic lines. Whiskers represent the limits of the 1.5 interquartile range. Letters indicate significant differences based on the results of Kruskal-Wallis one-way analysis of variance and post-hoc Fischer's LSD test ($p < 0.01$), n=18.

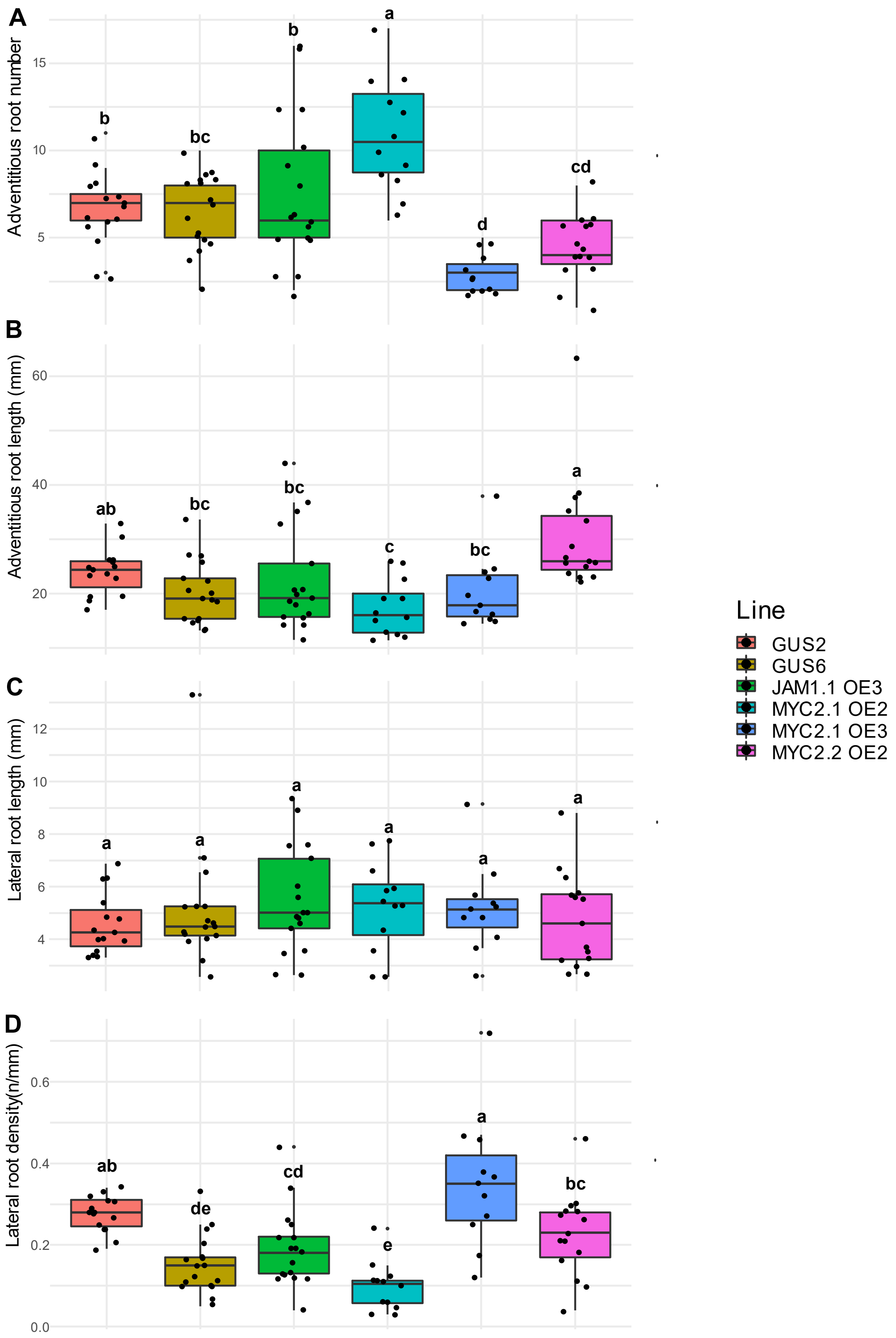


**Figure S3. The overexpression of *PtaMYC2.1, PtaMYC2.2* and *PtaJAM1.1* differentially alters root architecture.** Boxplots showing the distribution of **(A)** adventitious root number, **(B)** adventitious root length, **(C)** lateral root density and **(D)** lateral root length of transgenic lines. Whiskers represent the limits of the 1.5 interquartile range. Letters indicate significant differences based on the results of Kruskal-Wallis one-way analysis of variance and post-hoc Fischer's LSD test ($p < 0.01$), n=18.

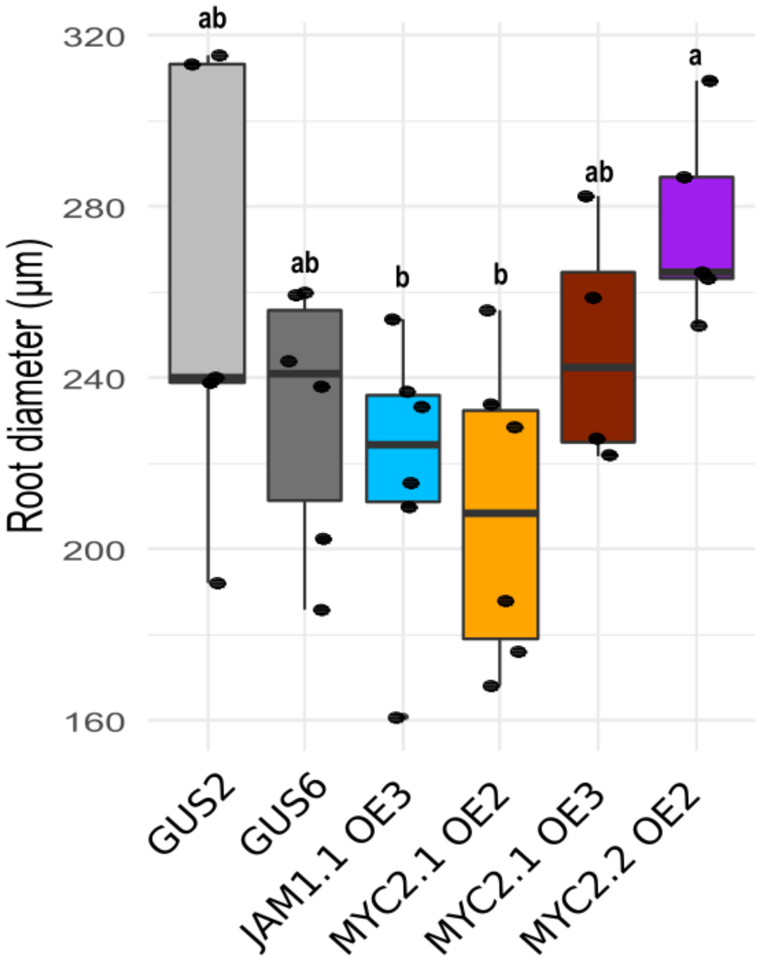


**Figure S4. The overexpression of *PtaMYC2.1, PtaMYC2.2* and *PtaJAM1.1* does not affect root diameter of ECM root tips.** Boxplots show root diameter of colonized transgenic lines. Whiskers represent the limits of the 1.5 interquartile range. Letters indicate significant differences based on the results of Kruskal-Wallis one-way analysis of variance and post-hoc Fischer's LSD test ($p < 0.01$), n=6.

>pMYC2.1 (1872bp)
CTATAACTAAGGTTGTAGGTTGACCGAGATCATAGTTATTAAACCCGGCCTAGGGATTGACCCGGTCAAGGGGCCGAGTCCCGGGTTTCATGGGCCAACTCGGGTCAACCCGGAAAAATTAAAAAAATATATTTAA
AGCTTTAATATTTTATATGAAAAAATTAAGAAAAAATCCATGTAAATATAAGCTATACATATTGTAAATAATGAAGTTTAAAAGAATATTTTAAAAAGTTTTTTATTCCATATTAAAAAGATATTGTGATAAACTTTGAAGTT
AAACTATTTAAACTAAATTTTTTTTATCCCACATTGAAGAAATAACGTTTTCATTGGGAACATAGAGTATATATAATAATGAGTTTCAAATCCCACATTGAAAAAACATAATTTTTTTTTATGGGAACAAAGAGTATACATA
CAAAAAGGCTTCAAATTCCACATTGAAAAGATATTATGTTATCCCTTTAAGTTGAAGTATTTAAACCAAAAACTTTTTTATCCCACATTGAAAAAACACAATTTTTTTCTTGGGAACATAAAGTATATATACTAATGGATT
TCAAATCTCACATTGAAAAAAACACAACTTTTTTCATGGAAACATATAGTATATATACGAAAGGGCTTCAAATCCTACATTGAAAAGATATTATGTTATCCCTTTAAGTTGAAGTATTTAAACCAAATTTTTTTTATCTCA
CATTGAAAAAACATAATTTTTTTTGAGAACATAAATTATACATACTAATGGATTTCAAATCCCACATTTGAAGAAAAAAACACTAGGTCTTGTTCGGGTTTGCCTGGGTCGACCTGGTCATGGTTGACCCGCTTGGTCG
AACGGGGTTTGCCGGGCTATTGCCTCGGCCAGTCTTTTATTAAATCCGAACTGGTTCAGCAACCGGGTCAACCCGCCGAGCTGCGCCAAGCCAGGTTTTATAACTAGGACCGAGATCAATCTAATTTGACGTCATC
ATAATATTAAAAAAATTATTATCTTAATTTTTTTTAAAAAACTATAGCTTATGTTTATCAATTATCTAAATTATCTTTCATCTTATTAAATTAAATAATTTATGTGCGTTTGGCATTATATTTCAAAAGTGCTTTTGAAAAAAT
TAAAAATTTTTTTTACTTTAAATTAATATGTTTTTGATGTTTTCAAATCATTTTAATATGCTGATGATTTTTAAAAAATAAAAAAATATTATTAACATACTTTTTTAAATAAAAAATATTTTAAAAATAATCACAACCACATTTC
TAACCCTATAAGCTGGATTTACAGATTTTAAATTGAATCACTTTATTAAATTTAATAATACTGCTGACAAAATTCTACTCACCTCAGTCAAAACACTATCTAAATACGAACAGAACTATGCCACGTAATACCCACGTGCT
TCCACTGCATAACAATAGGCTAACCCTAACTCAGTCTAGAGCTGTAGAGTGATATGACGTTAACGCCAGCCTTTCCCTTCCATTCTAAAATCTCGTGAATTAATCGGCGACCCTTTCCTTCCTAGAAACTGCCACCCT
CACTTTCCACTCCCGTTCTCAACTTCACTCTAAAAAATTTAAGCTCAAATTCATACCAAATAAATCCCAAACCCAAACGCACCCCAAGTTCTAGCTCACCTTCCTCTCCTGTCCTCCTCCTCCCCTCCTCCTCCTCCT
CCTCCTCCTCCTCCACCACCACCACCACCACCACCACAGCAACCCCTCTCTCCGAAACAAATCTCAACCAAAAACACCACTTCCTCTCTCTGAAATAAACTTATTTTCACCACTTTCTCCCTCGAAAACAAACTCTTT
AATCCCAGCCCTATCTCCCTTTGAAAAAAAAAAACACATCTGCTTTCTGGG

>pMYC2.2 (1668bp)
TACTAGATGATCCAGATTCATCAATAACACGACTGCCTCCTGCTCGTGCTTTTGCGTGAGTGACCTTATTTTCGGTCACGATATCATAAAAAAATTAAAATCTATCCAAAGTTTTCAAGTGCATCAGGACTACCAAGAA
ATTTTATAATTGATCTAGTAATGGATGACCACTAACATTCCACTTCTTCTATGATTGGTCTTCAATTTCTTTAATTCAATCTTCAATCATGTAATATACATCACAATTAATTACAAATATAACAAAAGTTAGCATTCCCCTT
AACAATTTTATATAAACTCTAAAATTTCTAAATTAAAAATTTTTCAATATGAAACAAAACAAGAAAAAAAGTGGAAAATAATACATTTCCAATGCTAATTAATTCAAATTGACATCCCTTTTTCAACAATTTCAAGAATCTC
CTAATTTTCTTCCAAACTCCAAACTTCCTCTCTCTAATCACCCAAGCTCTCCATCTCTCTCTCTAAGTTTTTTCTCTACTTTGATTTGGTTTTCTCTGCTTGGAAATCACCAAAAACCAATCTCATTTAATTGGGAGGAA
TTAAAGAAAAAGAAGAAGAATTAGAATAAATAAACAAGCTAATTTTGAGGGGTGTAACTCAACTGGTCAGGCTCTAGGTTTGCTCCCTAGAAATCACCAGTTCGAGTCTCACAAACCTCACGGTCACTGGAGGCTTA
TATAATCGTTAATTTAGGACCCGTGGGATTAGTCGAGGTGCACGCAAGTTGGGCCGGACATCCATATTAATAATAATAAAAAAAAGGCTTAATTCATAATTCACATGAAGATCATAACAAATAGATAATCTTGATTAAA
TTTAATTTAATGAGAGAAAAAGTGGTGATAAAAAGAATTAAAATTGAGTTTTAATTAGGACATGACGGAAAATCAATTGAAAAAGCAGATGAGAGGGAAATGTGAACAAACCTACATGGCAAGGAGTGAGAATCATAC
AAAGTATAAATGTGAACCTTTTTTCACATACTATATGGTGCATTTGATTATTTATTATTTAATCTCACACCTCTTGGTCCAACTCCTCATGGAGATGTAGCAATCATTTCAGTCCAAAGAGGAGTCACCGAGTCCAATAT
TAGGTAAGCTTCCATAAATACCCCCCACCAGCTGATTTCCAACTTCCTTCTCTACGAGTCACCTCGGTCCAAACCAGCAGCTCCAAACCAGCGGAACTATTCCACGTGATCCCCACGTGCTCCTATAGCCAGCCCT
AAACCCAGCCTTAACTGCATCCCCTCCAGTCCACGTTCACGCCGCCCTTTACCCTTCGCCCTTCAAAATATCGTGAATTAATCGGCGACCCTTTCCTTCCTAGAAACTGGCACCCTCACTTTCTTTCCAATCCTCTA
AATTAACTCTTTAAAAAATAATAATATGAAATTAGAAAAATTAAGCTCAAATTCACTCCAAATAAATTCCAAACCCAAACGCAACCCAGATTACTCCCCTCCCGCCCCTCCCCTCCACCACAACCACCACCACACGCTC
TCTCTCTGAACCAAATCCCCATCAAAACCACCGCTTTCTCTCTCCAAACTAAACTTCATTTCACCCCTTCCTCACTCGAAAACAAAAGTAAGGATCTCCCTTTATTTAAAAAAGAAACAACATATATTTACTTTTTGAG

>pJAM1.1 (1667bp)
ATATTTTTTTAAAAATTAATTTTTTTAATATATTTTGAATAGTTTTTCATTCAAATTAAAATCCTAACAAGCATGAGATAAGAAAAAGAAAGGAAGGAAGAACAAGACGAAGCAACAATTGGAAACGATTAGCAACGCTGT
CAGAATTTTTCCTTCCCACCTGCCGAGGGAAGTCAATGAGTTGCAGATATTTATTTCTAGTTAGATTTTTTCCGCATCCAACTCCAACTTCAATTCATTCATTCATCTTCCATGTCACTTCTCTCTATGACGATTGAGA
GTTTTTTTTATGTTTTTTTTCAAGTGGTTCTTACTCTAAAATATATTAAAATAATATATATTTTTTTTAAAAAAATATTTTTGATGTTAATAAATTAAAATAATAAAAAATATTTTTTTAAAATATCCCTGGATATCTCCCTTGCC
ACTGGATTTCAAACTTGAGGTCGGTTTATGGGGGCGTAAAATCCTCATAAGTGGAAAACAACAGATTTGTTTTTAACACATGGAAATTATTCTTTATAAATGAAAATAAAAAAGAGAAGAAATAATCATGCCATGCAAA
CCATGAAAAAGAATCCAAATTATGTGACGAGCAAATAAGTAGCGGGCCGCAAACAGTAAGAAAAACCAGTGGATCCCAATAATTTAGGACAATGGTTAACAAATGTGATTTGTGTTAAGAATGTAACAGAGGGGTGC
GGTTTGTTAAGAAAGAAGAGGAAGAAGAAGAACATAACAAGTGTTAAAGCTGATCGGATGTCCGCCACTGTTAACCAACTATGGAATCTAACCTTGATGTCTGGATAAAAAGATGGTGTGTAAACAAGGGTTTTTTTA
AAGGGTTTCCAGGTGAGCCTACTGTTTACTGCTTTTATTAGACCAACAAACAAGCTCCACAAAATCGTTGCCGACTATCTCTCAACCTCCAACAATTCTTTCTTTGTTTTTATTTTCATCTCTCAATCAAAATACTATTA
AATTTTAGGTTTCTTTTATTTAATCATGCATTTAACCATGGTTACTTTTTTTAATCCAATCTATACTTAACCATGGTCCACTTATGCAGTTAAGGACTAAGAGGGCAGAACACCAAACTGCAAATCCACTCCTCTCTCTC
TCTCTCTCTCTCTCTCTCTCATTTTCCACCCTTTGGCTCTAAAGAGAGAAAAAGCCTTTCCCTTCCTTTTCCCATTCCTTATTGTTTCTGAAATATTTAACAAATTTTCTGCTTGCTTTCCGGGATTTTTTTTTTTTATAAA
CCTGTCCCTTTACCTGGAAAACTTATACCACCTGAGAATAAATACCATCCTGCTACCTTACCCCAATTGTTGTCTCATTATCCCAATTTGCCTGACGCAACCGGTGTGTATTTTCTTTCAATTGTCTCTTCTTTGCATC
AATCTACTTTACTTTTTCTGGGTTTTCCTTGCAACTTGTTATTTTTGCTATTTTAACTTATTGAGTTCCTGAGATCTGAATTTATGCATATAGTATTGTACGAGAGAAAACATGTGGGCTTGTTTTATTTTGCTAAATGCA
TTAGATTTTAGCCCCTATCTATTATAGAAACTAGGGGGGTTCAGTACCAAGATTTGATTCTGTGTAATAAAGTTTGAAGCTTTGTGTTTAGTCGGTAGTTTGAAGTTTCAGTTTTGGGTGTCCTAGAA

**Figure S5. Promotor sequences of PtaMYC2.1, PtaMYC2.2 and PtaJAM1.1**

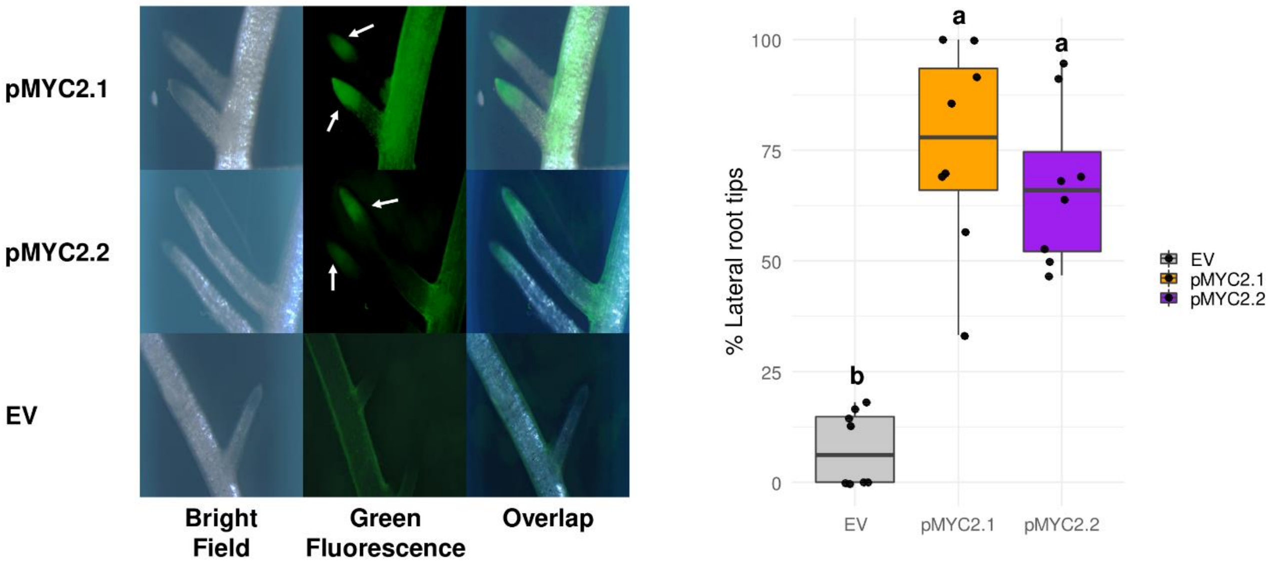


**Figure S6. PtaMYC2.1 and PtaMYC2.2 are both expressed in poplar lateral roots.** The left panel shows representative stereomicroscope images of poplar hairy roots transformed with pMYC2.1::GFP, pMYC2.2::GFP or empty vector (EV) under the bright field or with green filter (470 nm excitation, 525 nm emission). Boxplots in the right panel represent the percentage of lateral root tips displaying GFP fluorescence under stereomicroscope observation. Whiskers represent the limits of the 1.5 interquartile range. Letters indicate significant differences based on the results of Kruskal-Wallis one-way analysis of variance and post-hoc Fischer's LSD test ($p < 0.01$), n=8.

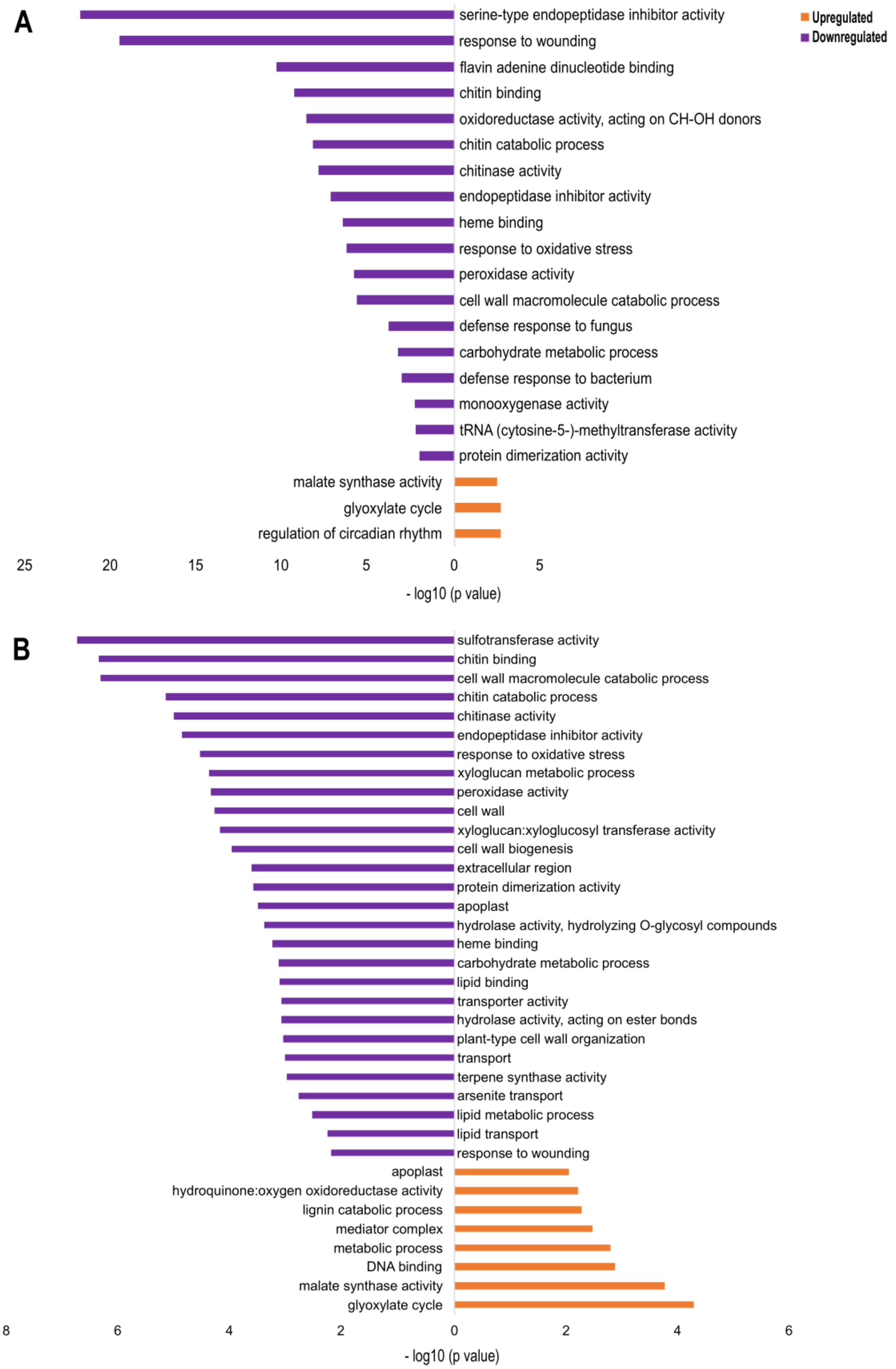


**Figure S7. Genes downregulated in the JAM1.1 OE line are enriched in defense functions.** Gene ontology (GO) enrichment was performed over **(A)** JAM1.1 OE3 vs GUS6 (Dataset S8) and **(B)** JAM1.1 OE3 ECM vs GUS6 ECM (Dataset S9) gene sets. Analyses were made separately for up and downregulated genes. Only significantly enriched ($p < 0.01$) terms are shown.

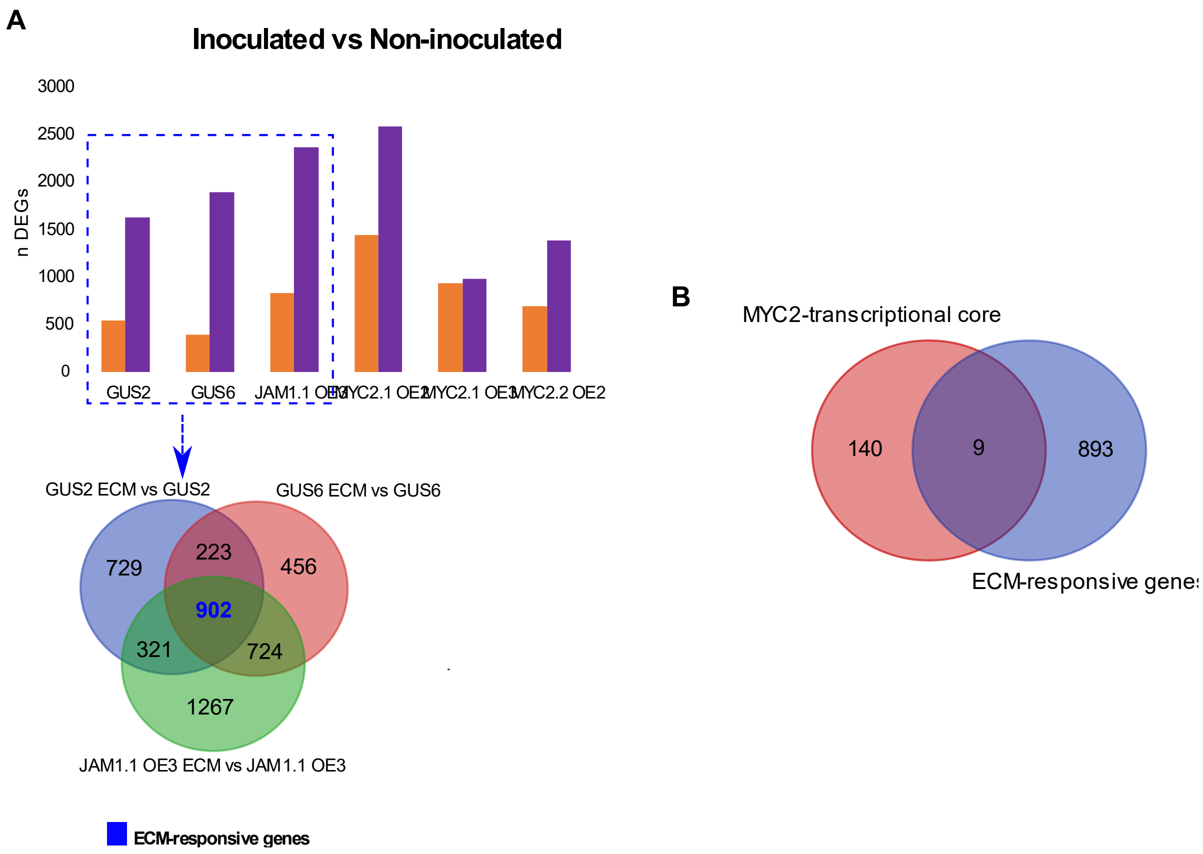


**Figure S8. Differentially expressed genes in response to ECM formation in JAM1.1 OE3, GUS2 and GUS6 lines. (A)** Number of differentially expressed genes (nDEGs) in RNA-seq data fromECM 21 days post infection (dpi) and non-inoculated roots (inoculated vs non-inoculated) for each transgenic line. Venn's diagram representing the "ECM-responsive gene set" formed by 902 DEGs common to the lines showing proper ECM structures (GUS2, GUS6 and JAM1.1 OE3) in inoculated conditions vs non-inoculated conditions. **(B)** Overlapping genes between MYC2-transcriptional core and ECM-responsive genes represented in a Venn's diagram.

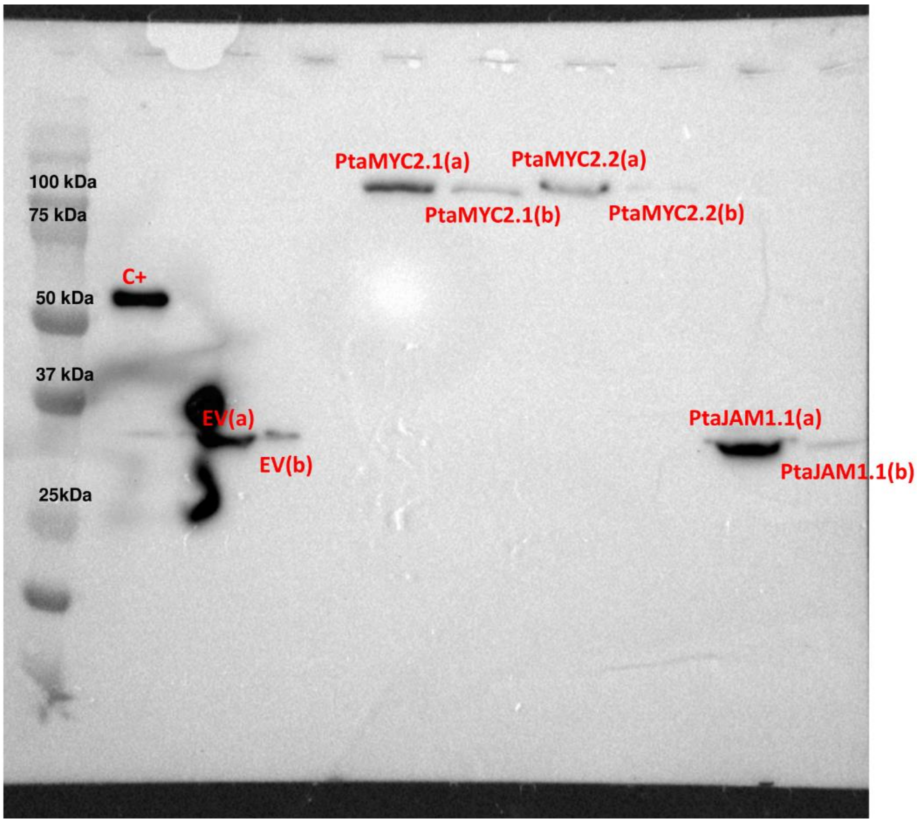


**Figure S9. *In vitro* expression of pIXHALO Empty Vector, PtaMYC2.1, PtaMYC2.2 and PtaJAM1.1.** Western blotting was performed using anti-Halo Tag monoclonal antibodies (Supp. Materials and methods). From left to right, the lines were loaded with Page ruler protein ladder, HaloTag® standard protein (Promega, C+), Empty vector (EV), PtaMYC2.1, PtaMYC2.2 and PtaJAM1.1. (a) Refers to the *in vitro* expressed protein before Halo bead binding and (b) is the washed protein after Halo bead binding.

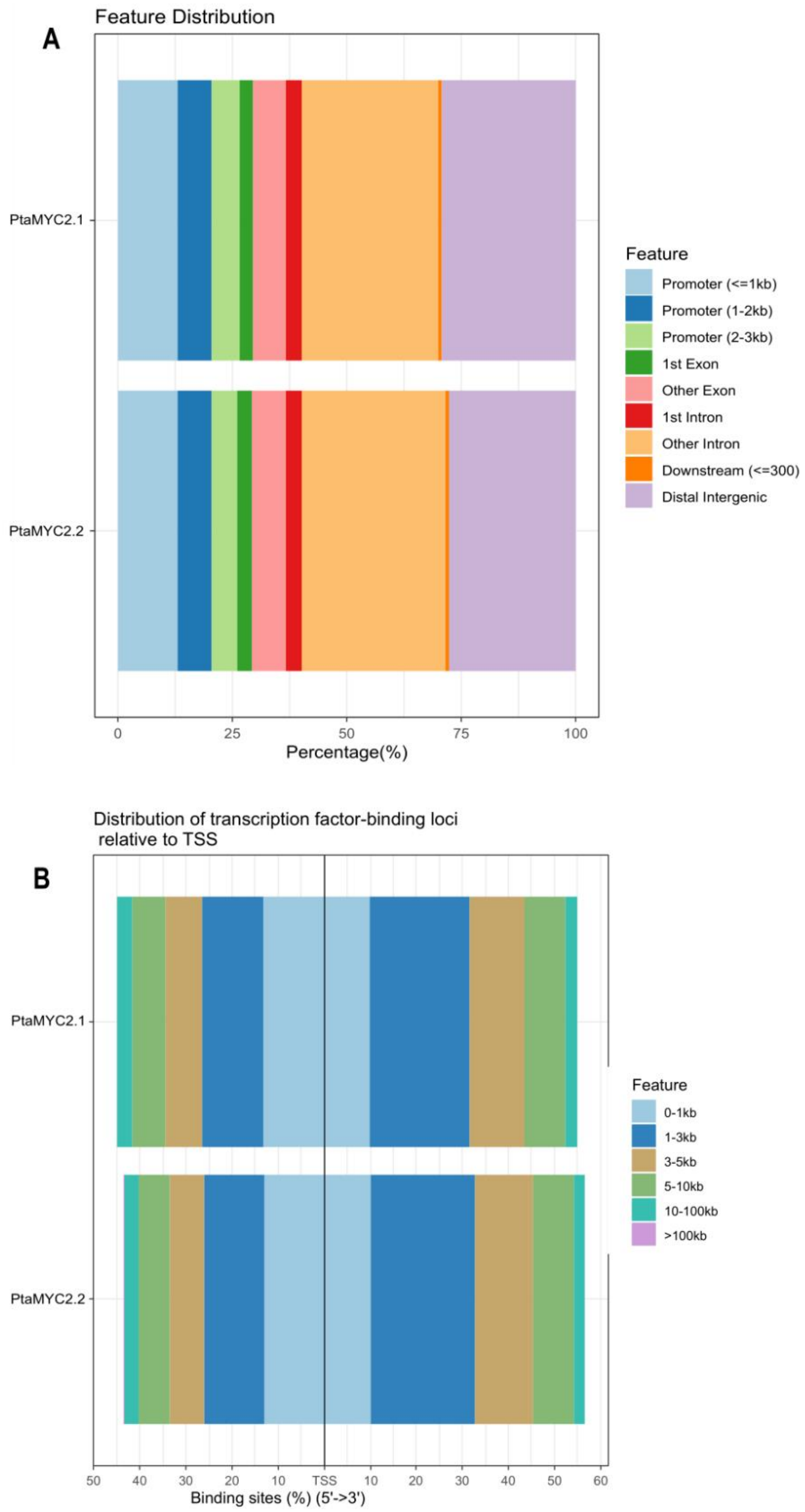


**Figure S10. Distribution of PtaMYC2.1 and PtaMYC2.2 DAPseq peaks according to their genomic features. (A)** Percentage of each feature in the total peaks and **(B)** distribution of peaks according to their relative distance to transcription starting sites (TSSs).

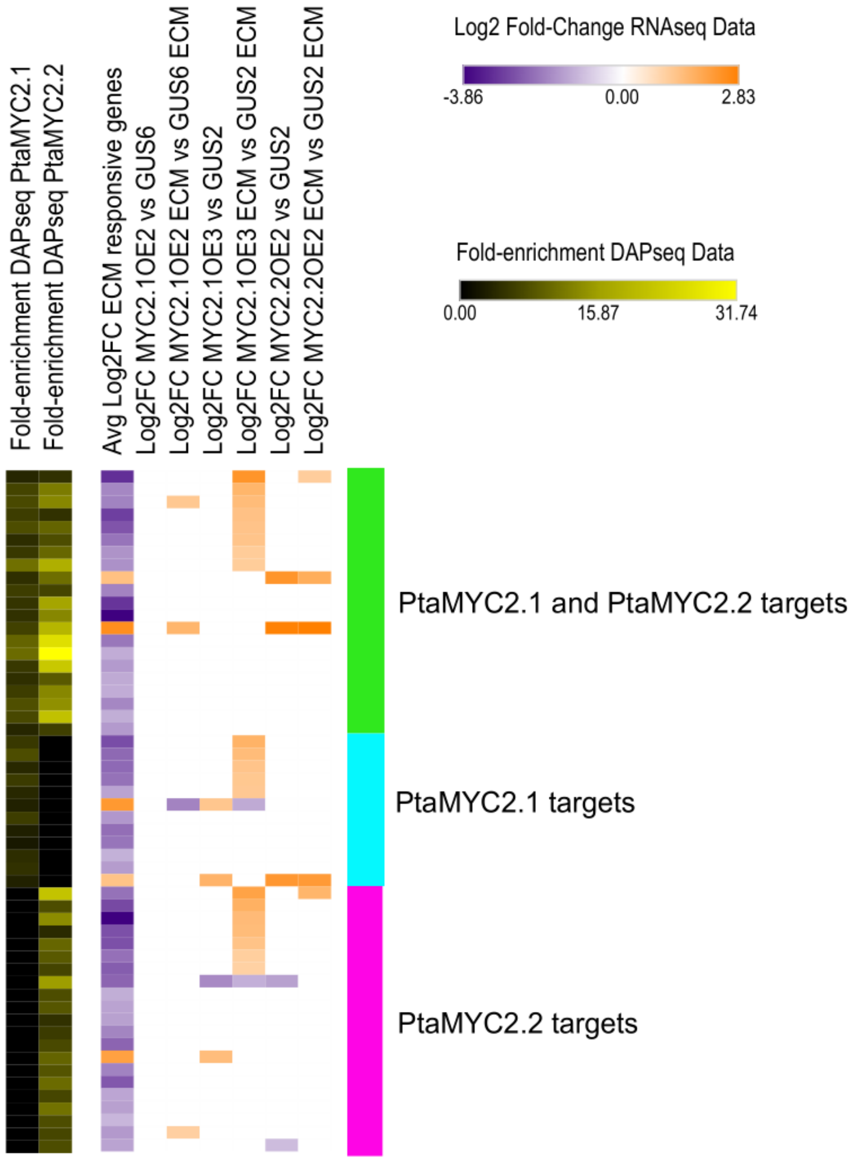


**Figure S11. Several ECM responsive genes are directly targeted by PtaMYC2.1 and PtaMYC2.2 TFs and upregulated in MYC2.1 OE3 ECM.** Heatmap showing the DAP-seq peak fold-enrichment (left) for the 54 genes overlapping between the DAP-seq data and the RNA-seq ECM responsive genes. A second heatmap (right) shows their expression profiles according to RNA-seq data. The average log2 fold-change of GUS2, GUS6 and JAM1.1OE3 in inoculated vs non inoculated conditions, and the log2 foldchange of MYC2 OE lines vs their control lines in both ECM and non-inoculated conditions are shown.

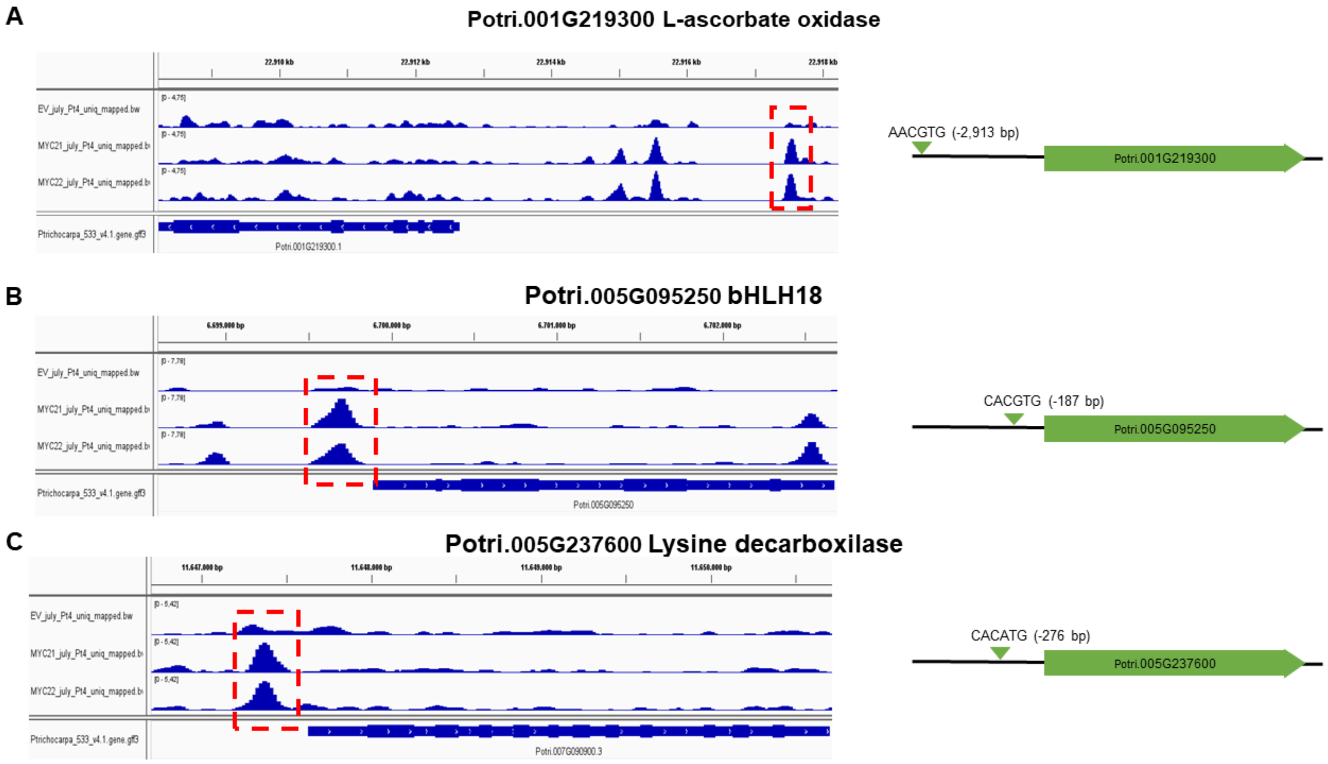

A
Potri.001G219300 L-ascorbate oxidase
AACGTG (-2,913 bp)
Potri.001G219300
B
Potri.005G095250 bHLH18
CACGTG (-187 bp)
Potri.005G095250
C
Potri.005G237600 Lysine decarboxilase
CACATG (-276 bp)
Potri.005G237600

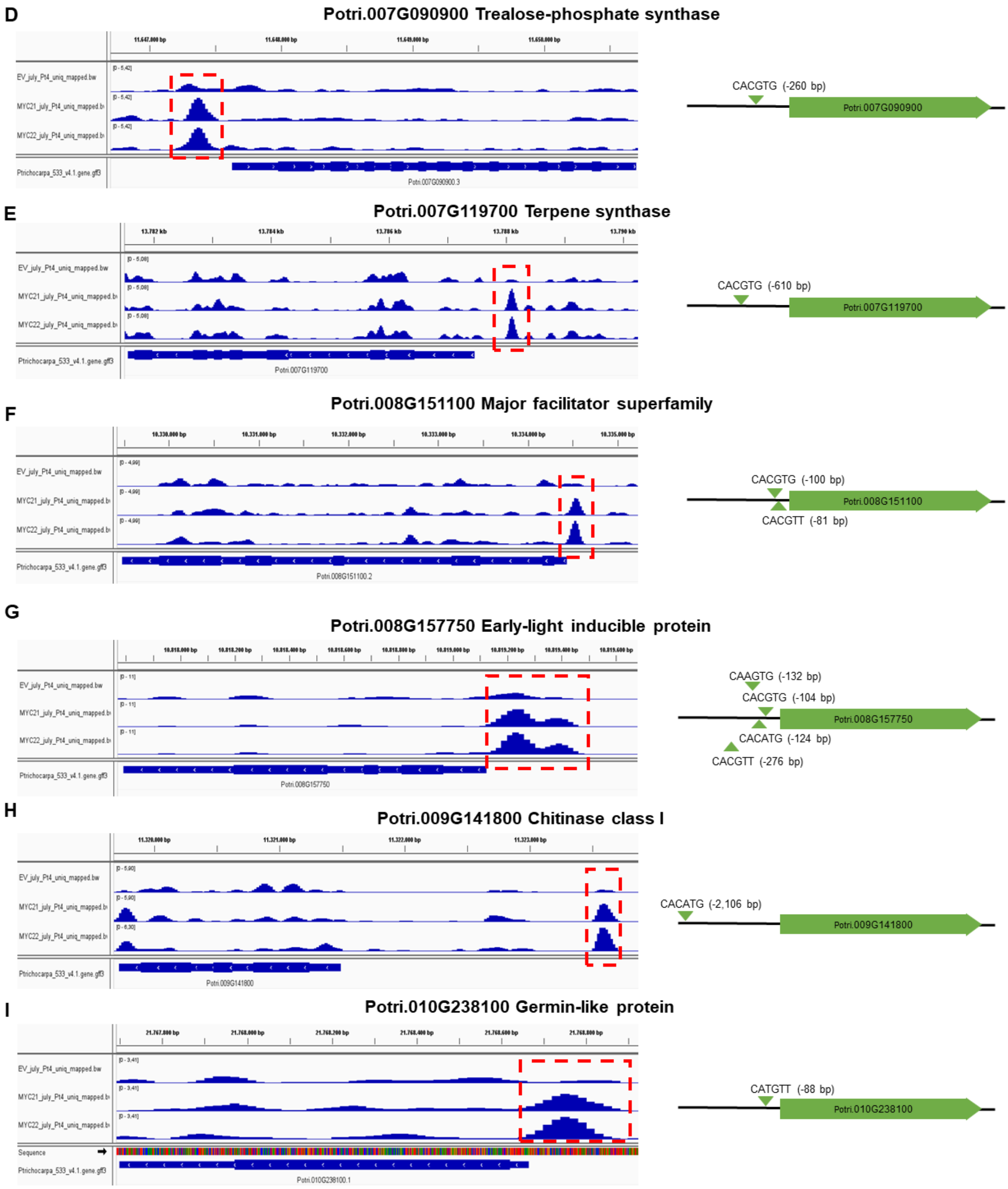
D
Potri.007G090900 Trealose-phosphate synthase
EV_july_Pt4_uniq_mapped.bw
MYC21_july_Pt4_uniq_mapped.b
MYC22_july_Pt4_uniq_mapped.b
Ptrichocarpa_533_v4.1.gene.gff3
Potri.007G090900.3
CACGTG (-260 bp)
Potri.007G090900
E
Potri.007G119700 Terpene synthase
EV_july_Pt4_uniq_mapped.bw
MYC21_july_Pt4_uniq_mapped.b
MYC22_july_Pt4_uniq_mapped.b
Ptrichocarpa_533_v4.1.gene.gff3
Potri.007G119700
CACGTG (-610 bp)
Potri.007G119700
F
Potri.008G151100 Major facilitator superfamily
EV_july_Pt4_uniq_mapped.bw
MYC21_july_Pt4_uniq_mapped.b
MYC22_july_Pt4_uniq_mapped.b
Ptrichocarpa_533_v4.1.gene.gff3
Potri.008G151100.2
CACGTG (-100 bp)
Potri.008G151100
CACGTT (-81 bp)
G
Potri.008G157750 Early-light inducible protein
EV_july_Pt4_uniq_mapped.bw
MYC21_july_Pt4_uniq_mapped.b
MYC22_july_Pt4_uniq_mapped.b
Ptrichocarpa_533_v4.1.gene.gff3
Potri.008G157750
CAAGTG (-132 bp)
CACGTG (-104 bp)
Potri.008G157750
CACATG (-124 bp)
CACGTT (-276 bp)
H
Potri.009G141800 Chitinase class I
EV_july_Pt4_uniq_mapped.bw
MYC21_july_Pt4_uniq_mapped.b
MYC22_july_Pt4_uniq_mapped.b
Ptrichocarpa_533_v4.1.gene.gff3
Potri.009G141800
CACATG (-2,106 bp)
Potri.009G141800
I
Potri.010G238100 Germin-like protein
EV_july_Pt4_uniq_mapped.bw
MYC21_july_Pt4_uniq_mapped.b
MYC22_july_Pt4_uniq_mapped.b
Sequence
Ptrichocarpa_533_v4.1.gene.gff3
Potri.010G238100.1
CATGTT (-88 bp)
Potri.010G238100

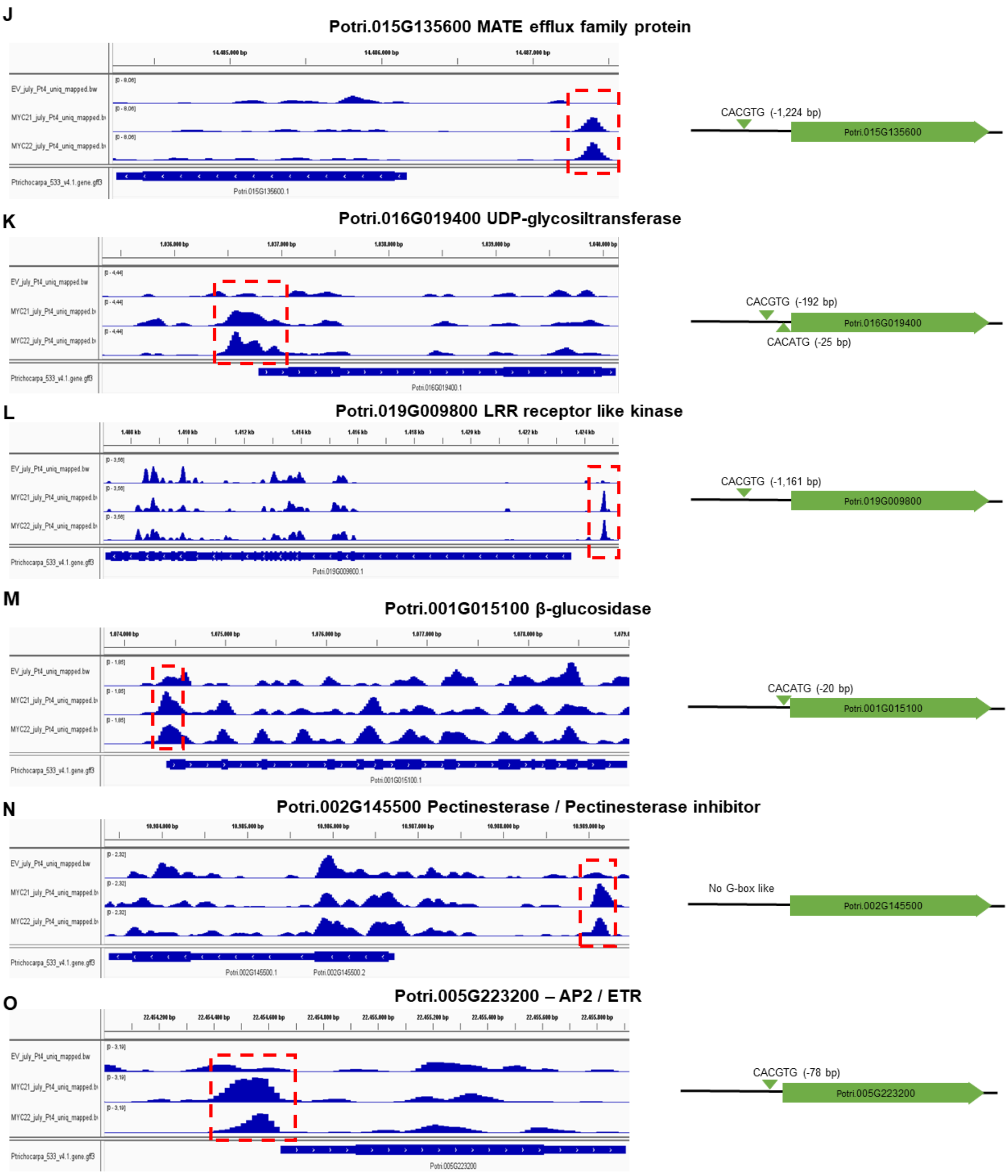
J
Potri.015G135600 MATE efflux family protein
CACGTG (-1,224 bp)
Potri.015G135600
K
Potri.016G019400 UDP-glycosiltransferase
CACGTG (-192 bp)
CACATG (-25 bp)
Potri.016G019400
L
Potri.019G009800 LRR receptor like kinase
CACGTG (-1,161 bp)
Potri.019G009800
M
Potri.001G015100 β-glucosidase
CACATG (-20 bp)
Potri.001G015100
N
Potri.002G145500 Pectinesterase / Pectinesterase inhibitor
No G-box like
Potri.002G145500
O
Potri.005G223200 – AP2 / ETR
CACGTG (-78 bp)
Potri.005G223200
EV_july_Pt4_uniq_mapped.bw
MYC21_july_Pt4_uniq_mapped.bw
MYC22_july_Pt4_uniq_mapped.bw
Ptrichocarpa_533_v4.1.gene.gff3

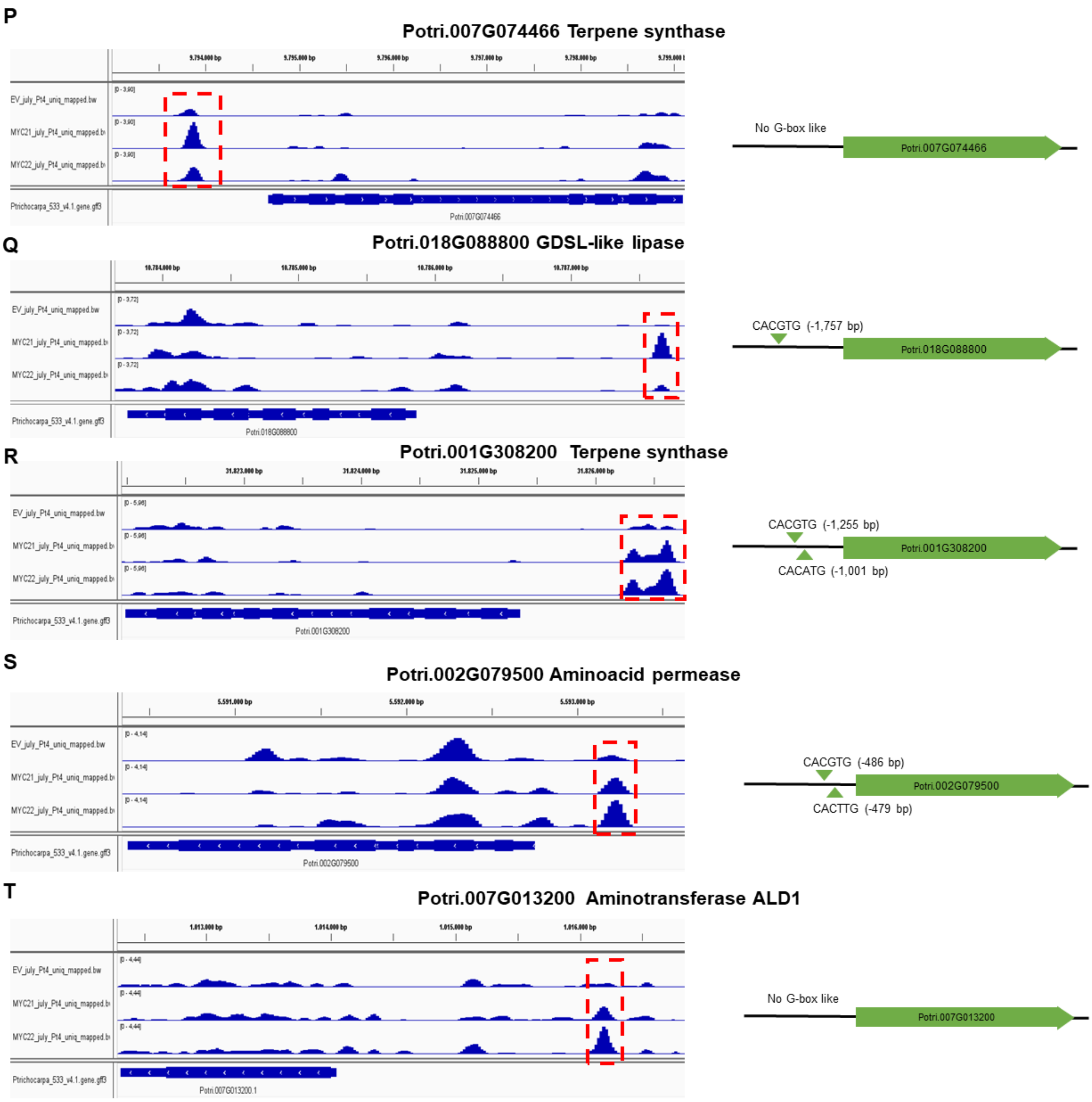


**Figure S12. IGV browser view of DAP-seq peaks adjacent to MYC2 transcriptional core genes.** Every gene **(A-T)** is represented with a blue box and its orientation is indicated by the direction of the white arrow inside the blue box. MACS2 peaks (blue peaks) denote either PtaMYC2.1, PtaMYC2.2 or Empty vector (EV) DNA binding sites (DBSs). Significantly enriched peaks are highlighted with a red dotted box. To the right, a schematic representation of the genomics coordinates of G-box like motifs found near to the summits of each significant peak is presented.

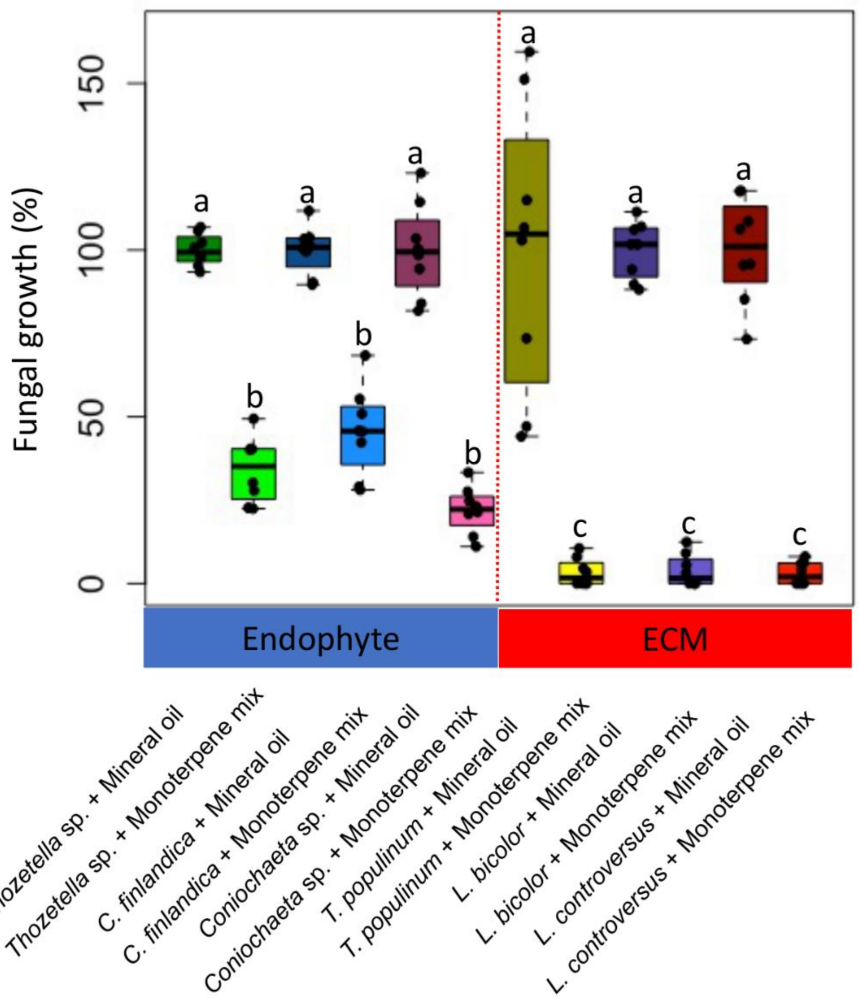


**Figure S13. Monoterpenes produced by PtTPS16 and PtTPS21 inhibit fungal growth of different ECM and endophyte species.** Boxplots showing *in vitro* fungal growth of each species in presence of mineral oil 0.5 µL $cm^{-3}$ (control) or monoterpene mix 0.5 µL $cm^{-3}$ (γ-terpinene, (-)limonene, (-) camphene, (-)α-pinene and (-)β-pinene, 0.1 µL $cm^{-3}$ each). Whiskers represent the limits of the 1.5 interquartile range. Letters indicate significant differences based on the results of Kruskal-Wallis one-way analysis of variance and post-hoc Fischer's LSD test ($p < 0.01$), n=8.

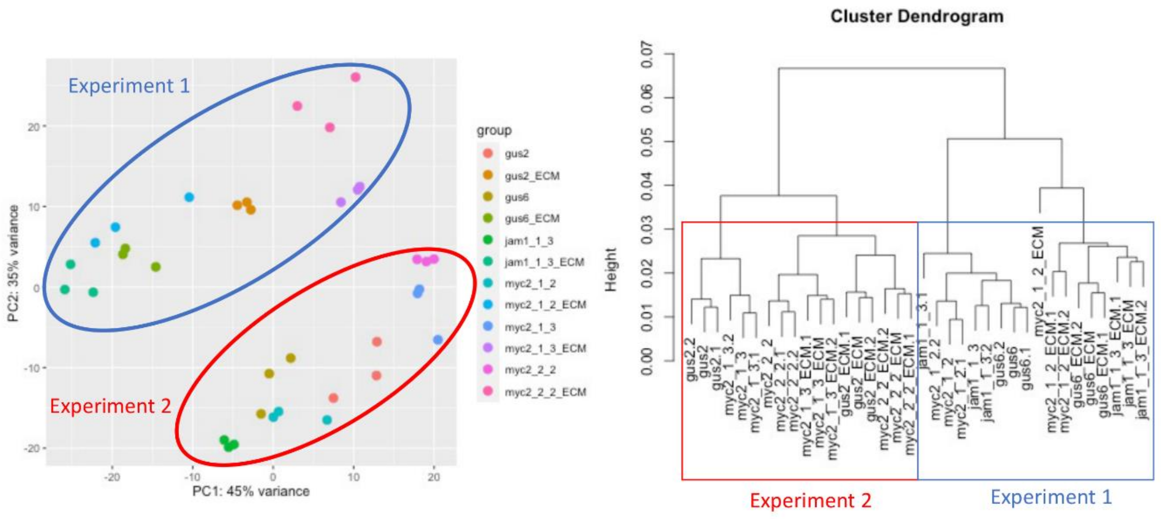


**Figure S14. RNA-seq genome-wide data clusters according to our experimental setup.** Principal component analysis (PCA, left) and hierarchical clustering (right) of the *vst* normalized transcriptomic data using DESeq2 R package.
